# Comparative genomics and phylogeny of sequenced IncHI plasmids

**DOI:** 10.1101/334409

**Authors:** Senén Álvarez

**Keywords:** Plasmids, IncHI, Tra Regions, Multiple Drug Resistant (MDR) Modules, Phylogeny

## Abstract

Conjugative plasmids from the IncHI incompatibility group are relevant vectors for multidrug resistance. Is the objective of this article the screening of the phylogeny of the first 44 publicly available sequences of IncHI plasmids from Enterobacteriaceae (1961-2013), because since there, a few hundred were registered in Genebank. A comparison of the similarities and differences between them revealed extensive conservation of their coding sequences. However, each genome had accessory regions with features of degradation or resistance to antibacterial agents. This article elucidates the IncHI1, IncHI2 and IncHI3 subgroup plasmids from the evolutive line based on multiple drug-resistant modules, explains the origin of the IncHI1 pAKU1-like composite transposon lineage, typifies the subgroup IncHI3, which is distantly related to IncHI1 and IncHI2 in phylogenetic analyses, confirms the relations within the transfer regions of H-group plasmids, and points to the divergence time of evolution with phylogenetic trees. Homology between subgroups, as measured by the molecular clock test, indicated that the contemporary IncHI1 and IncHI2 transfer region lineages diverged from an ancestral clone approximately 8,000 years ago (17,000 years ago if the divergent IncHI3 plasmids are included), although the IncH backbone was older. The uniformity of the composite transposon shared by most IncHI1 subgroup plasmids might be functionally equivalent to the heavy metal resistance that is mediated by some operons in the IncHI2 subgroup plasmids. A comparison of the DNA among these closely related plasmids provided insights on the horizontal and vertical transfers that were involved in their development.

## 1. INTRODUCTION

Bacterial plasmids have a wide range of properties, including genes mediating resistance to various antibiotics. Plasmids are classified into incompatibility groups according to their inability to be maintained within the same host cell. Plasmids belonging to the H complex by classical incompatibility tests have been classified into two groups: IncHI1 and IncHI2 (Whiteley and Taylor 1983), which were increased, by genetic testing, with the IncHI3 group (Subramanian et al. 2016).

The common regions of the IncHI plasmids contain genes that are essential for plasmid replication, maintenance, and transmission (which might define the general backbone for the group), as well as an extensive series of homologous core genes. Acquired DNA, that is not common among plasmids core, includes class 1 integrons, transposons, insertion sequence (IS) elements, and genes for other phenotypes that are advantageous for survival, such as resistance to antibiotics, toxic heavy metals, colicins, and bacteriophages (Gilmour et al. 2004; Johnson et al. 2006

Bacterial conjugation spreads DNA throughout populations. Conjugation is an efficient mechanism for genetic exchange among bacteria, and it requires three plasmid-encoded multiprotein complexes: a membrane-spanning mating pair formation (MPF) complex (which encodes determinants for transfer), the mobilization (MOB) machinery (which encodes a complete set of transfer genes and the cytoplasmic nucleoprotein relaxosome complex), and the relaxase or the homomultimeric type IV coupling protein (an inner membrane ATPase that is the conduit for DNA transfer and the determinant of the specificity and functionality of the conjugative pore). (Alonso et al. 2005; Alvarez-Martinez and Christie 2009; de la Cruz et al. 2010; Guglielmini et al. 2013; Wong et al. 2012). All of these genes are located in two regions of the IncHI plasmids (called the Tra regions), which sharing determinants of the MPF and MOB mechanisms (Alonso et al. 2005; Lawley et al. 2002; Lawley et al. 2003b; Rooker et al. 1999).

Plasmids, as members of the bacterial mobile gene pool, are important contributors to horizontal gene transfer (HGT) between bacteria. HGT contributes to bacterial evolution and adaptation by the acquisition of radically new genetic information, expanding genomic evolutionary processes beyond the pre-existing genes (Dominges et al. 2012; Fricke et al. 2009). Hence, knowing the precise locations of the resistance genes enables the identification of related plasmids and facilitates the monitoring of how they spread (Cain and Hall 2012b; Cain and Hall 2013).

The aim of this work was to leverage the availability of the DNA sequences of single bacterial plasmids by looking for phylogenetic relationships within the IncHI plasmids, according to the characteristic original core of the group and with the genetic modules acquire as consequence of the evolutionary pressure. The IncHI plasmids studied were sequenced from the first one characterized (R27, isolated in 1961), until those isolated in 2013; since then, the IncHI sequenced plasmids are several hundred.

## 2. MATERIALS AND METHODS

### 2.1. IncHI plasmids

The 44 plasmids analysed in this study were sequenced before June 2015. Their general characteristics (name, date of isolation, country, original bacteria, length, GenBank entry, and references) are in Supplementary Data 1a: “IncHIPlasmids”. The annotations of the complete prokaryotic IncHI plasmid sequences were obtained from GenBank (http://www.ncbi.nlm.nih.gov/genbank/). Three plasmids (the pAKU1-like plasmids p9804, p7467 and p6979) were sequenced by the Wellcome Trust Sanger Institute and were downloaded as FASTA files (http://www.sanger.ac.uk/resources/downloads/bacteria/salmonella.html> 454 Sequencing >Illumina Data).

IncHI plasmids were detected using a concatenated Tra genes sequence, representative from each taxon or clade, with BLASTn in GenBank. Concatenated Tra genes sequences for detection of new sequenced plasmids are in Supplementary Data 1b: “IncHI plasmids Concatenated Tra genes”. Many other IncHI plasmids, not included in analysis, were sequenced since June 2015; some comments and relation with previous are mentioned in results.

### 2.2. Comparative sequence analysis

Unless otherwise stated, sequence comparisons at the amino acid or nucleotide level used the BLAST algorithms from the National Center for Biotechnology Information (http://www.ncbi.nlm.nih.gov/guide/all/#tools). Multiple sequence alignment used ClustalW2 from the European Molecular Biology Laboratory-European Bioinformatics Institute (http://www.ebi.ac.uk/Tools). Other tools from these sites (Uniprot, MUSCLE, EMBOSS, etc.) were also used. Plasmid phylogeny was determined using the BEAST v1.8 package (http://beast.bio.ed.ac.uk/) and theMEGA5 program (www.megasoftware.net). The accession numbers for other genetic elements with homologues to the partial IncHI plasmid sequences are in Supplementary Data 1c: “GenBank Data”.

### 2.3. Molecular clock test

The complete Tra1 and Tra2 regions of the IncHI plasmids were downloaded from Genbank database and protein-coding gene sequences were extracted. The selected Tra1 genes were *trhH, trhG, trhRY, traG, traI* and the homologous genes from the plasmids at the R27 loci R0115, R0116 and R0118. The selected Tra2 genes were *trhK, trhC, parA, eexB, parM, trhU, trhN, trhI* and the homologous genes from the plasmids at the R27 locus R0009. Equivalent analysis was developed for the IncHI1-pNDM-CIT-like, IncHI2 and IncHI3 plasmids to the TerZF/TerWY operon (*terF, terE, terD, terC, terB, terA, terZ, terW, terY, terX, terY2, terY3*). The H-group plasmid chosen were pRA1 (IncA/C), pSN254 (IncA/C), R391 (IncJ-ICE), SXT (ICE), Rts1 (IncT) and pCAR1 (IncP7); the transfer genes selected for analysis were *traJ, traG, traI, trhF, trhH, trhG, trhU, trhW, trhF, trhC, trhV, htdT, trhB, trhK* and *trhE*.

Using the same Tra genes from the IncHI plasmids (excluding the plasmid pCFSAN002050-2, which lacks the transfer regions, pEC-IMP similar to pEC-IMPQ, and pEQ1 similar to pEQ2), ClustalW alignment (according to coding direction) was performed for homology comparison and utilization as concatenated supergenes. Distance estimation (dS = number of synonymous substitutions per synonymous site), was determined using the BEAST and MEGA programs for displaying and printing molecular phylogenies, molecular clock hypothesis and ages of divergence (Holt 2009; Zhang et al. 2006).

To calibrate and fit ages of divergence for the IncHI plasmids, are used data for ten sequenced IncHI1 plasmids (excluding pNDM-CIT-like plasmids, which belonged to a divergent lineage) isolated between 1961 (R27) and 2008 (pEQ2). The phylogeny of the IncHI1plasmids has been studied (Phan et al. 2009; Holt et al. 2011) for an acceptable calibration of a synonymous substitution rate for this time period (Duchêne et al. 2016; Salichos and Rokas 2013).

The ages of divergence were calculated for each taxon using the formula AGE= dS x (Years of analogous pool/ dS of analogous pool), which corresponds to AGE= dS x 47/5,514×10^−4^ = dS x 85,237 (dS length data were obtained from the expanded branch of FIGURE1). With this formula for the closely related IncHI1 Tra multisequences (which evolved in a short time), objections to the mutation rates (MR) and generations per year (GY), which involve copy number, are avoided.

The ages of divergence are calculated as well using a theoretical formula: AGE = dS/(2 x MR x GY) years (Pearson et al. 2009), where MR is the mutation rate and GY is the generations per year (Drake 1991; Ochman et al. 1999). In this case, using suitable determination with IncHI1-R27-like (dS=5,514×10^−4^, MR=5,5 x 10^−10^, and GY=300) gave a result of AGE = 5,514×10^−4^/(2 x 300 x 5,5 x 10^−10^)=1671 years, rather than the actual 47 years. To adapt these dates, a calibration factor (CF) between the hypothetical and the actual values is necessary (CF=47/1671). With this CF, the formula can be expressed as AGE=2,8126 ×10^−2^ x (dS/3,300×10^−10^) = 85,232 x dS, which is the value of the simplified formula (AGE=85,237xdS). Additionally, the 14 sequenced IncHI1 R27-like plasmids come from *S*.*enterica*, except for three from *Escherichia coli* (pECO111, pEQ1 and pEQ2) and one from *S*.paratyphi (pAKU1), which are not distant enough phylogenetically to assume a relevant divergence in MR and GY.

Finally, the years calculated with this formulation and mentioned in results are the consequence of the determination hypothesis.

## 3. RESULTS AND DISCUSSION

### 3.1. IncHI PLASMID HOMOLOGY

The sequenced IncHI plasmids shared the common structural composition of an extended series of core genes that were equivalent in each subgroup and nearly identical between plasmids of each subgroup. Sequence genes of each, are in Supplementary Data 2: “Homologous Sequences”, arranged by homology alignment.

In general, excluding the divergent plasmids, approximately 130 genes were shared among the studied IncHI, or 160genes if we include those that were deleted in some plasmids, or the genes that were not identified as coding sequences

The common genes in the IncHI1, IncHI2 and IncHI3 subgroups were the transfer genes required for conjugative functions and the genes implicated in copy number control, maintenance, ultraviolet light protection, and thermoregulation of conjugation, among others. Nevertheless, some features were shared by only some of the plasmids in each subgroup and were either lost or not acquired by the others.

A variety of genes and operons (Supplementary Data 2: “Homologous Sequences”), had interesting relationships to the phylogenetic differentiation of the IncHI1 plasmids, and two subgroups were detected: R27-like plasmids and pNDM-CIT-like plasmids. (References where IncHI1 plasmids have been described before are: Akiba 2007; Dolejska et al. 2011; Dolejska et al. 2012; Dolejska et al. 2013; Holt et al. 2007; Kubasova T et al., 2016; Ogura et al. 2009; Ong et al. 2012; Parkhill et al. 2001; Sherburne et al. 2000).

In subgroup IncHI2, the core differences are not significant between the plasmids. The acquisition of MDR determinants deleted the adjacent open reading frames and occasional sporadic insertions occurred with some genes and transposases. Nevertheless, the homogenous characteristics of the IncHI2 plasmids core in respect to other subgroups are obvious. The case of plasmids pCFSAN002050-1 and pCFSAN002050-2, both isolated in a multistate outbreak of *Salmonella enterica* subsp. *enterica* serovar Cubana infections linked to alfalfa sprouts, is interesting. Both plasmids together form a complete IncHI2 plasmid sequence, which means that the original IncHI2 plasmid was broken in their ancestor, and both segments remains in the isolated strain. (References where IncHI2 plasmids have been described before are: Chen et al. 2007; Chen et al. 2009; Conlan S et al., 2014; Feaseye NA et al., 2014; Gilmour et al. 2004; Han et al. 2012; Hoffmann M et al., 2014; Johnson et al. 2006; Kariuki S et al., 2015; McGann P et al., 2015; Sheppard AE et al., 2015; Timme et al. 2013).

Subgroup IncHI3 was unique. Gene correspondence in the core was sequential for some determinants; as occurs with genes from the previous section until the end of the Tra2 region, within the Tra1 region, and in the tellurite operon. In the rest of the core, some determinants were in order, but many others were displaced, and some were duplicated. Sequences that were not shared by rest of IncHI studied plasmids were seen, but they were homologous to other plasmids and genomes. According to the CDSs of the core, the two subgroups are clearly different (IncHI3-pNDM-MAR-like and IncHI3-pKOX-R1-like). (References where IncHI3 plasmids have been described before are: Becker L et al., 2015; Doi Y et al., 2014; Huang et al. 2012; Li J et al., 2014; Liao et al. 2012; Pereira PI et al., 2014; Schluter A et al., 2014; Stoesser N et al., 2014; Villa et al. 2012).

Recently, Liang et al (2017) proposed, based in similar results, a scheme for typing IncHI plasmids into five separately clustering subgroups, which includes the three mentioned for taxa, IncHI4 for clade pNDM-CIT, and IncHI5 for clade pKOX-R1. Nevertheless IncHI1 and IncHI4 belong to same taxon, as occurs with IncHI3 and IncHI5.

### 3.2. IncHI PLASMID BACKBONE

Plasmid transfer requires the expression of plasmid-encoded transfer genes (Supplementary Data 3: “Backbone”). Curiously, Tra regions have the same location in all sequenced IncHI plasmids, indicating that a precursor plasmid had this characteristic and that their separation (if they were previously together, as was hypothesised) occurs before. However, in H-group plasmids, the Tra regions are distant as well, with a pool of genes in each region. Which suggest, if both Tra regions were together, that was before of diversification from H-Group.

#### 3.2.1. Molecular clock test

The results obtained with the general formula (AGE= dS x 85,237) can only be used for guidance. The main objection is that the correlation between years and SNP variation needs the introduction of a statistical formulation because all of the IncHI1 plasmids, except for R27 (isolated in 1961), were collected in 24 years. As the divergence is quite similar for both pools (with or without R27, which means 47/dS_a_ or 24/dS_b_), the final AGE that is determined is one half. To understand this discordance, a similar determination was made with the pAKU1-like plasmids (pAKU1, p9804, p7467 and p6979; isolated in three years from Pakistan, Nepal, Morocco and Central Africa). In these plasmids, the shared composite transposon is presumably deactivated as a consequence of partition and inversion (Holt et al. 2007). This result suggests that these plasmids had only been evolving for the length of time between their isolation dates and that their ancestor is the same plasmid with the same composite transposon. The ratio years/dS for these four pAKU1-like plasmids was nearly 60% (3/9,369×10^−5^), which is lower than the IncHI1 determination, and suggests that determinations with 24-year pools could be more than adequate for study.

The clock rate (μ) of different bacterial lineages varies from 9×10^−9^ to 3×10^−5^ mutations per nucleotide per year (Achtman 2012; Duchêne et al. 2016), but no studies were done with plasmids of the accepted variability. With the obtained dS from the R27-like plasmids Tra genes sequence, (and not with the expanded FIGURE 1, to prevent divergences), the clock rates (μ=dS/years) can be calculated (Supplementary Data 4: Tra Genes Sub-Tree 1). As all of the studied IncHI1 R27-like plasmids have a value of dS=4,821×10^−4^, the corresponding clock rate for 47 years is μ=1,02 x 10^−5^ and for 24 years is μ=2,0 x 10^−5^. The same determination for the pAKU1-like plasmids, which were isolated in three years, was μ=2,07 x 10^−5^ and the result for the pHCM1-like pool was μ=2,04 x 10^−5^. These results are homogeneous to fix the clock rates for the IncHI plasmids. Although Duchêne et al. (2016) estimated that scale data can vary by more than an order of magnitude in sets less than 10 years, these clock rates have an average of nearly μ=2×10^−5^. Finally, a convenient formula for calibration and posterior extrapolation is AGE= dS x 24/5,514×10^−4^ = dS x 43526. (The option chosen, according to 24 years, is more feasible than the previous calculation to 47 years. In addition, it is more adequate to comment on and remember years rather than dS).

**FIGURE 1.**
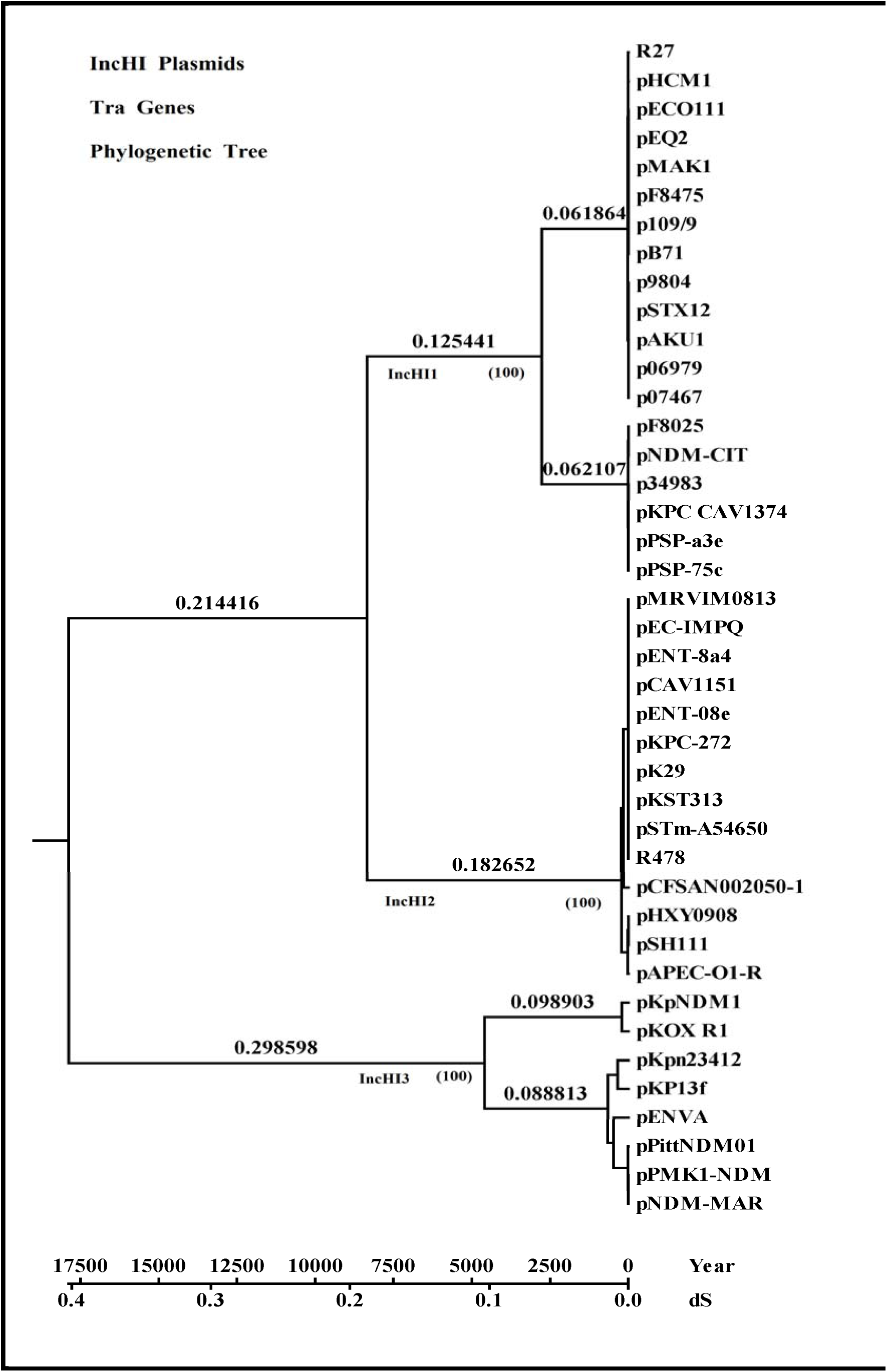
Evolutionary relationships within the Tra genes of the IncHI1, IncHI2 and IncHI3 taxa. Correspondence between synonymous substitutions and the years of divergence is shown. The tree is drawn to scale with branch length in the same units as the evolutionary distances used to infer the phylogenetic tree. Evolutionary analyses were performed using BEAST. The tree was drawn with MEGA5, using the Newick file. Bootstrap percentages, in parentheses, indicate the reliability of the cluster descending from that node and were tested with same file in MEGA 5.

The phylogenetic homologues of the concatenated *Tra* genes are in the phylogenetic tree (FIGURE 1). Four plasmids (pP-stx-12, pKpNDM1, pKP13f and pENVA) have deficient Tra regions (lacking some of the genes), but BEAST analysis without these deficient plasmids displays a similar tree with a 1% higher divergence. Three well characterized taxa are defined by the tree: IncHI1, IncHI2 and IncHI3. Using the previous considerations, the estimated complete divergence from a common ancestor corresponds to 17,509 years ago (dS=0,40228), whereas the contemporary IncHI backbone of the IncHI1 and IncHI2 subgroups diverged from an ancestral clone approximately 8,176 years ago.

To clarify the evolution of plasmids from every taxon or clade, the phylogenetic trees of each subgroup were displayed in the Tra gene tree (FIGURE 1), using the Newick file from BEAST and the MEGA5 program. The subgroup trees are in Supplementary Data4: “Tra genes Sub-Tree”.

Two clades are displayed by the IncHI1 plasmids, with a divergence time of 1,953 years. The clades correspond to the R27-like and pNDM-CIT-like plasmids, which suggest a parallel evolution in different ecological niches (Poirel et al. 2011). Both clades and the same subclusters were identified by Kubasova et al. (2016) using core backbone regions, and Liang et al (2017) using *traI* and *trhC* genes.

In the IncHI1-R27-like subtree (Supplementary Data4: Partial Tree 1b), there is a group of plasmids (pP-stx-12, pAKU1 and the homologuesp07467, p9804, p06979) that is different from the rest. The CDSs core differences in this group confirm divergence. This entire group shares the citrate operon, Tn*6026* (*betU*), except for p07467, which has lost the MDR modules as well. All of these plasmids share the same location of Tn*10* (with *tetB*, for tetracycline resistance).They conserve (except for pP-stx-12) the inversion of an extensive sequence that includes the Tra2 region, Tn*5393* (*strAB*), and the composite transposon that is shared by the other IncHI1 plasmids. The relations between the remaining plasmids from clade R27-like (Liang subtype IncHI1), as the subtree reflects, are not as coherent. The presence or absence of the magnesium/nickel/cobalt transporter gene (*corA*), citrate operon, composite transposon, or particular CDSs explains their phylogeny (Supplementary Material 2:”Homologous Sequences”). What is relevant is that this R27-like clade is the only one that does not have the Ter operon (for tellurite resistance). Since June-2015 to June-2018, 17 new plasmids has been sequenced of this clade (99% of identities with pHCM1 Tra genes); one of them (pKV7; GB: LT795503), is very similar in core and MDR to pB71).

Clade IncHI1-pNDM-CIT-like plasmids (Liang subtype IncHI4), show a very similar core genome and a uniform *Tra* gene phylogenetic tree (Supplementary Data4: Partial Tree 2). The BEAST divergence time respect R27-like clade is 2,705 years. The divergence time between plasmid of this clade was 13,5 years (they were isolates between 1990 and 2013). Homology from a common ancestor between pNDM-CIT and pF8025 is proven by their shared Tn*2501*/Tn*2502* (arsenical operon) and Phage CP4-57-like segment. However, differences in the MDR modules reflect the 20-year period between isolations. The rest of the clade plasmids (pPSP-75c, pPSP-a3e, pKPC_CAV1374) do not have differences in their core sequences prior to some insertions or deletions of punctual CDSs. Nine new plasmids have been sequenced since June-2015, with 99% of identities with pNDM-CIT. One of them (pLC476; GB: KY320277), is similar to pF8025 (with the Phage CP4-57-like segment from pNDM-CIT plasmid, and a different antibiotic resistance module).

Three clades can be recognized in the IncHI2 Tra genes phylogenetic tree (Supplementary Data 4: Partial Tree 3), with a divergence time of 226 years. The core sequences of the fourteen plasmids in the group are similar with respect to the insertion and deletion of CDSs, without any significant relation between them, even in clades. Their MDR modules are located in three places: near the Cop operon, beside the Kla operon, and adjacent to the Tn*1696* location. The most relevant characteristic is that the MDR modules are quite different (apart from some homology). Curiously, in the majority of plasmids, a module replaces the Cop operon-Sil operon-Tn*7* section, but this triple operon is shared by plasmids in each clade. Up of 100 new IncHI2 plasmids were detected, with BLASTn in GenBank, since June-2015 onwards; all of them display an identity of 99% with R478 Tra genes. Even if new clades are presents, divergence time will not be very different from the already mentioned. Three of those new sequenced plasmids were studied by Liang et al (2017) and by Sun et al (2016) and genetic conservation was observed within the core sequences of representatives IncHI2 plasmids, but differences in MDR modules; the comparison will be made in the corresponding section (3.3.6).

The third taxon of the IncHI plasmids is the IncHI3 taxon. Its phylogenetic distance with respect to IncHI1 and IncHI2 is reflected not only by the homologies between ORFs (frequently not detectable with BLASTn, but confirmed with BLASTp) but also by the displacement of many common CDSs to different position. This taxon, as occurs with the IncHI1 and IncHI2 taxa, has many CDSs that are only present in the IncHI3 plasmids (Supplementary Data 2: “Homologous Sequences”). Additionally, five CDSs of these plasmids are shared with the IncHI1 plasmids and four CDSs are shared with the IncHI2 plasmids, which indicate that this taxon comes from a common ancestral plasmid with CDSs remaining in IncHI1 and IncHI2. As in IncHI1-CIT-like and IncHI2, the IncHI3 plasmids have the TerZF/TerWY operon, which suggests that the common ancestral IncHI plasmid probably had the TerZF/TerWY operon. Two clades (Supplementary Data 4: Partial Tree 4), with a distance estimation of 4,512 years between the plasmids of each, were detected. Both show similar order in their sequences, but many genes were gained or lost in each clade. Beyond the hypothetical proteins, relevant differences among clades are the partial deletion of the TerWY operon and the insertion of the Tn*10* transposon in IncHI3-pKOX-R1-like plasmids (Liang subtype IncHI4).The MDR modules differ in their positions and sequences, but a common module is located after TerWY, in which the Mer operon (from Tn*1696* or Tn*21*) and other sequences, more or less diminished, are present.

Eleven plasmids were sequenced since June-2015 belonging to clade pKOX-R1; nine of them were isolated in *Klebsiella*, and two in *Raoultella ornithinolytica*. Identities with Tra genes were 99%, except for one, with 96% (p704SK6_1; GB: CP022144). One of them, (pYNKP001-dfrA; GB: KY270853), studied by Liang et al (2017), will be compared in the corresponding section (3.5.7).

The concatenated sequence of pNDM-MAR Tra genes (Liang subtype IncHI3), with BLASTn, display 35 new plasmids isolated in *Klebsiella* (except one in *E*.*coli* and other in *Pluralibacter gergoviae*, an Enterobacter). As eight plasmids display 98% of identities, and three 94% of identities, these conjectured new branches need to be confirmed in future studies.

### 3.3. IncHI1 PLASMIDS AND MULTIDRUG RESISTANCE

#### 3.3.1. MULTIDRUG RESISTANCE MEDIATED BY IncHI1 PLASMIDS

Analysis of the regions containing multiple antibiotic resistance genes that were inserted into the sequenced IncHI1 plasmids indicated that the insertions were distributed in the conserved IncHI1 core (Supplementary Data 2: “Homologous Sequences”).The two clades of this group (R27-like and pNDM-CIT-like) show different patterns of MDR modules, without any relation between them, but with homology between members of each clade.

The accumulation of resistance since the 1960s, when R27 was isolated, has been hypothesized as a consequence of the independent acquisition of resistance genes by a R27-like lineage since all plasmids share a related, extended backbone (Holt et al. 2011). However, careful study yielded confusing results. Before commenting on these results, it must be mentioned that the MDR modules and the determinants encoded by IncHI1 R27-like plasmids are in Supplementary Data5: “IncHI1 R27-like MDR” and the profiles of antibiotic resistance and the enzymes encoded by the resistance genes are in Supplementary Data 6: “Antibiotic Determinants”.

#### 3.3.2. PHYLOGENETIC RELATIONSHIPS AMONG IncHI1 R27-LIKE PLASMIDS

Analysis of the multiple antibiotic resistance gene elements shared by the R27-like plasmids indicated that the insertions were related.

##### 3.3.2.1. Transposon Tn*10*

All R27-like plasmids, except for p7467, encoded the tetracycline resistance transposon Tn10 with a complete sequence in R27, pP-stx-12, pAKU1 and homologues p9804 and p6979. The rest of the plasmids (pECO111, pHCM1, pEQ1/pEQ2, pMAK1, pB71, p109/9 and pF8475) had truncated sequences. Three different locations were observed, initially suggesting that the Tn*10* insertion was independently acquired in each plasmid rather than originating from a common ancestor (Gilmour et al. 2004; Holt et al. 2007). Careful study yielded several interesting results. (Supplementary Data 5: “IncHI1 R27-like MDR”).

The identical insertion in pP-stx-12 and pAKU1 pointed to a common ancestor, although pP-stx-12 did not have the multiresistance composite transposon. In pMAK1, pEQ1/pEQ2, pECO111, pB71, p109/9 and pF8475, Tn*10* was in a position adjacent to the IS1 genes from a multiresistance composite transposon. Duplication of Tn*10*-IS10 L in pHCM1 is a consequence of inversion. In addition, Tn*10* in these seven plasmids was truncated to IS10R and TetD, on the side adjacent to the composite transposon (Wain et al. 2003). In pP-stx-12 and the pAKU1-like plasmids, a complete Tn*10* was separated by four ORFs in respect to the rest of the above mentioned plasmids; although this result was remarkable, it might not be by chance.

Phylogenetic analysis of the IncHI1 plasmids was determined with common genes for Tn*10* using BEAST. Only three plasmids displayed SNP variations; as consequence, the tree did not have analytical significance.

##### 3.3.2.2. Composite Transposon

The similarity among the resistance gene complements of pAKU1, pECO111, pMAK1, pEQ1/pEQ2, pHCM1, pB71, p109/9 and pF8475 suggested the acquisition of a single composite transposon by these plasmids, with different rearrangements in each (Supplementary Material 5: “IncHI1 MDR”). The most relevant reorganizations are the inversion in pHCM1, the partial inversion in pMAK1, the location of two segments inside the Tra2 region in pAKU1 and derivatives, and the absence of the Tn*6029* and Mer operon in pB71 (Supplementary Data 2: “Homologous Sequences”).

Specifically, Holt et al. (2007) hypothesized that some plasmid precursors first acquired Tn*9*, followed by the transposition of Tn21 into Tn*9*. This point to Tn*2670*, which consists of an IS1-flanked segment containing the Tn*9*-like transposon that carries an In2 within Tn21 (Cain and Hall, 2012a), and found in plasmid NR1. At some moment, Tn*6029* was inserted into the integron In2 of Tn*21*, adjacent to *tni*. Consequently, the resulting composite transposon was propagated, with additional resistance genes inserted, and the deletion of In2 from Tn*21*. Once inserted into the other plasmids, the composite transposon was disrupted by new insertion elements. Thus, the most likely hypothesis was that the insertions to form a composite transposon occurred once and then moved as a single unit, using the IS1 ends of Tn*9* between plasmids (FIGURE 2 and more details in Supplementary Data 5: “IncHI1 R27-like MDR”).

**FIGURE 2.**
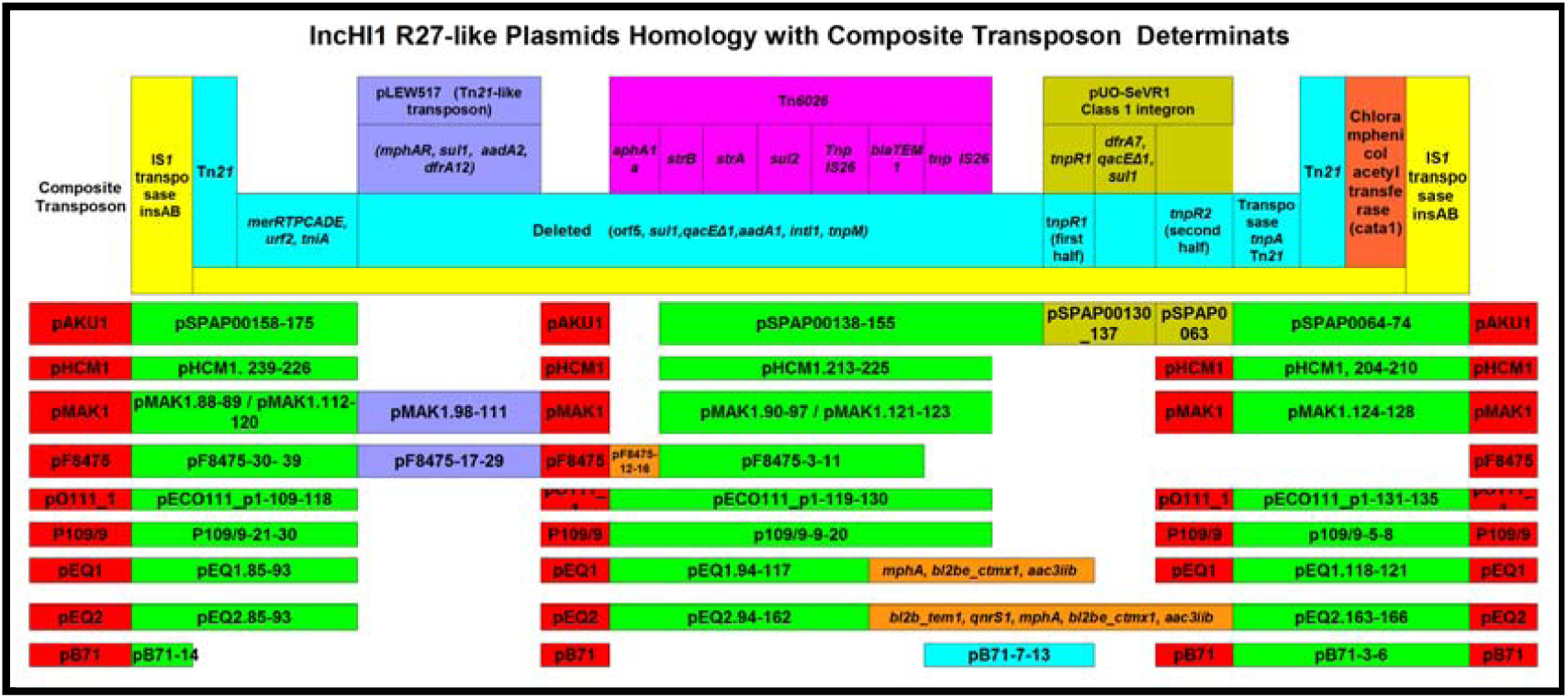
IncHI1 R27-like plasmid homology with composite transposon determinants. Green colour refers to the initial composite transposon. Dark blue, brown and orange refer to later additions. (Genes correspondence in Supplementary Data 2: “Homologous Sequences” and Supplementary Data 5: “IncHI1 R27-like MDR”).

In pAKU1, two IS26 insertions mediated two inversions in the composite transposon with a problematic difference (Phan et al. 2009). Adjacent to and continuous with *tnpA* was the first half of *tnpR* from Tn*21*; after an inversion from insertion of a class 1 integron pUO-SeVR1-like inside *tnpR* (Rodríguez et al. 2011),the second half of *tnpR* continued with *tnpM* and *intI1* from Tn21. In contrast, pHCM1-like composite transposon has not this *tnpR*. The presence of this *tnpR* suggests divergence between composite transposon lineages. However, a derivative of the pAKU1-like composite transposon (identified as a genomic island in an *S*.*typhy* that was isolated in Bangladesh in 2007 and studied by Chiou et al. (2015) highlights the problem, as it displays 100% identity for the complete sequence with an intact and non-interrupted Tn*21* resolvase gene.

The consequences of the pUO-SeVR1-like insertion are shown in FIGURE 3, in which the *tnpR* from Tn*21* comes from pUO-SeVR1-like.

**FIGURE 3.**
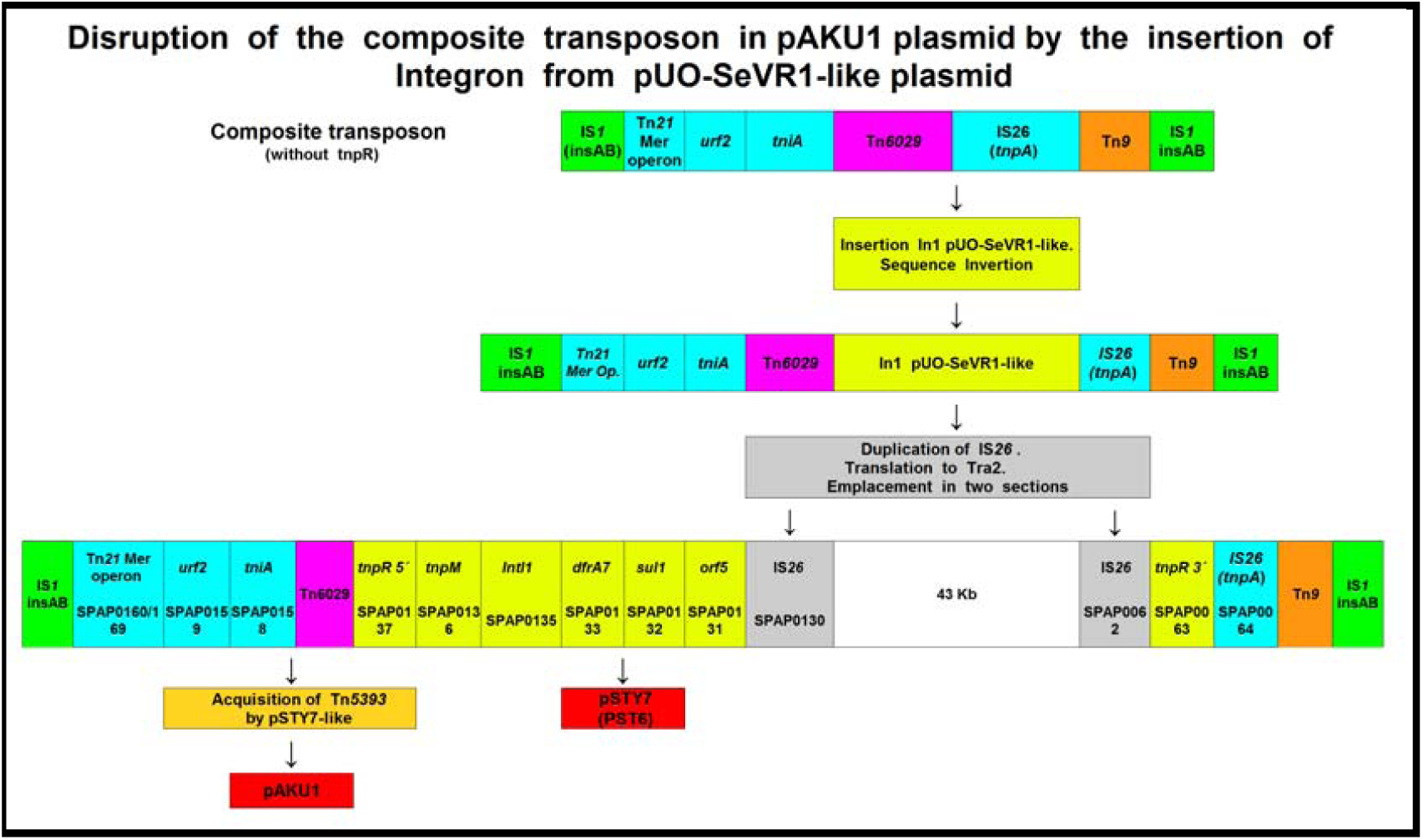
Hypothetical disruption of the composite transposon (IS1-Mer operon-Tn*6029*-Tn*9*-IS1) in a pAKU1-like precursor plasmid. The insertion of the pUO-SeVR1-like integron caused the rearrangement in the pAKU1 plasmid, mediating the inversion of the adjacent ORFs and their translocation to the Tra2 region from the composite transposon.

Another question is the location of the multitransposon in the pAKU1-like precursor before the rearrangement (in rest of plasmids it replaces the citrate operon). Nevertheless, pP-stx-12 had Tn*6062* (BetU), IS1 in the pAKU1 location is adjacent to the composite transposon, Tn*10* is in the identical position, a complete citrate operon is present, there is no inversion of the Tra2 region, and a single extended sequence (bp. 153742 to bp. 153941) that was truncate in pAKU1 (SPAP0257) and derivatives, but shared by other IncHI1 plasmids, is present. This suggests that pAKU1 and pP-stx-12 had a common ancestor and that the insertion of the multitransposon, with or without pUO-SeVR1, by horizontal gene transfer, caused their divergence.

pB71 is a particular plasmid in this situation. Isolated before the appearance of the composite transposon, it shares the citrate operon (deleted in rest of the subtype), the truncate Tn*10* is in an appropriate position, and don’t shows the composite transposon Tn*6029* neither the Tn*21* Mer operon (but the initial Tn*21* In2 sequence and the last IS1); which point, but not confirms, Tn*2670* origin. As this plasmid was the first IncHI1plasmid collected after R27 (in 1984-85; the composite transposon was detected during the same years in plasmid p109/9 from the same location; Faldynova et al. 2003 and Hradecka et al. 2008), a precursor of the pHCM1-like composite transposon can be considered. In other words, the later insertion of Tn*6029* and the Tn*21* Mer operon rearranged the composite transposon and deleted the citrate operon and the first half of Tn*21*. This hypothesis is feasible with the pHCM1-like plasmids, but it is inconsistent with the pAKU1-like plasmids because their Tn*10* transposon is located in a few distant CDSs. However, this initial Tn*21* segment in pB71 justifies the Holt and Phan hypothesis on the origin of the composite transposon (Holt et al. 2007; Phan et al. 2009). Probably pAKU-like plasmids received the complete composite transposon.

The plasmid pKV7 (GB: LT795503), sequenced as fasta file in 2017 (99,95 % of identities with pB71 Tra concatenated genes), is similar in core, exactly in MDR module (Tn9-In2 from Tn21-Tn10) and share, in another location, the complete sil operon-cop operon-Tn7, from plasmid R478 (with 99,96 % of identities).

##### 3.3.2.3. Relationship between composite transposon subtypes

Ten IncHI1 plasmids (isolated in the 1970s and early 1980s from various geographic sources) were studied by Taylor et al.(1989) to learn about the multidrug-resistance pattern. Phenotypic characterization revealed that all plasmids had chloramphenicol, streptomycin, sulphonamide and tetracycline resistance, one plasmid encoded as well ampicillin resistance, and one had trimethoprim resistance (resistance genes were not characterized). DNA homology was demonstrated with the *tetA* gene, but *tetCD* and the mercury and gentamicin resistance gene (*aph3Ia*) were not studied.

More recently, Holt et al. (2011) studied the resistance genes and insertion sites of 13 IncHI1 plasmids that were isolated before 1995 and showed different haplotypes with a significant relationship. The plasmids that were isolated in Asia between 1972 and 1977 did not have *tetD* from Tn*10* or the Mer operon but had a CatA1 determinant. In contrast, Latin American isolates between 1972 and 1981 had a complete Tn*10* and, in some cases, they also had the Mer operon and the CatA1 determinant.

These results about pHCM1-like (Type 2) and pAKU1-like (Type 1) and the partial homology of their resistance determinants might be coincidental, but it is still remarkable, since both subtypes were probably present at least ten years after the isolation of R27.To combine these subtypes, therefore, would require the acceptance of some unsupported speculations, such as the displacement of R27’s Tn*10* to the pP-stx-12/pAKU1 location, the joining of Tn*10* to the composite transposon in the pHCM1-like subgroup, and the acquisition of composite transposons by a pAKU1-like precursor plasmid, as pP-stx-12-like plasmid.

Tra gene analysis of the IncHI1 R27-like plasmids yielded nucleotide differences that were not remarkable in the diversification of the subtypes from a common ancestor. To clarify the relationships between plasmids, an 85-kb sequence of common DNA was extracted from the FASTA files of R27, pHCM1, pAKU1 and pP-stx-12. The sequences were concatenated in order, aligned with ClustalW2 and studied with MEGA5 software to determine the base substitutions between sequences with a Minimal Evolution Tree. The results are shown in FIGURE 4.

**FIGURE 4.**
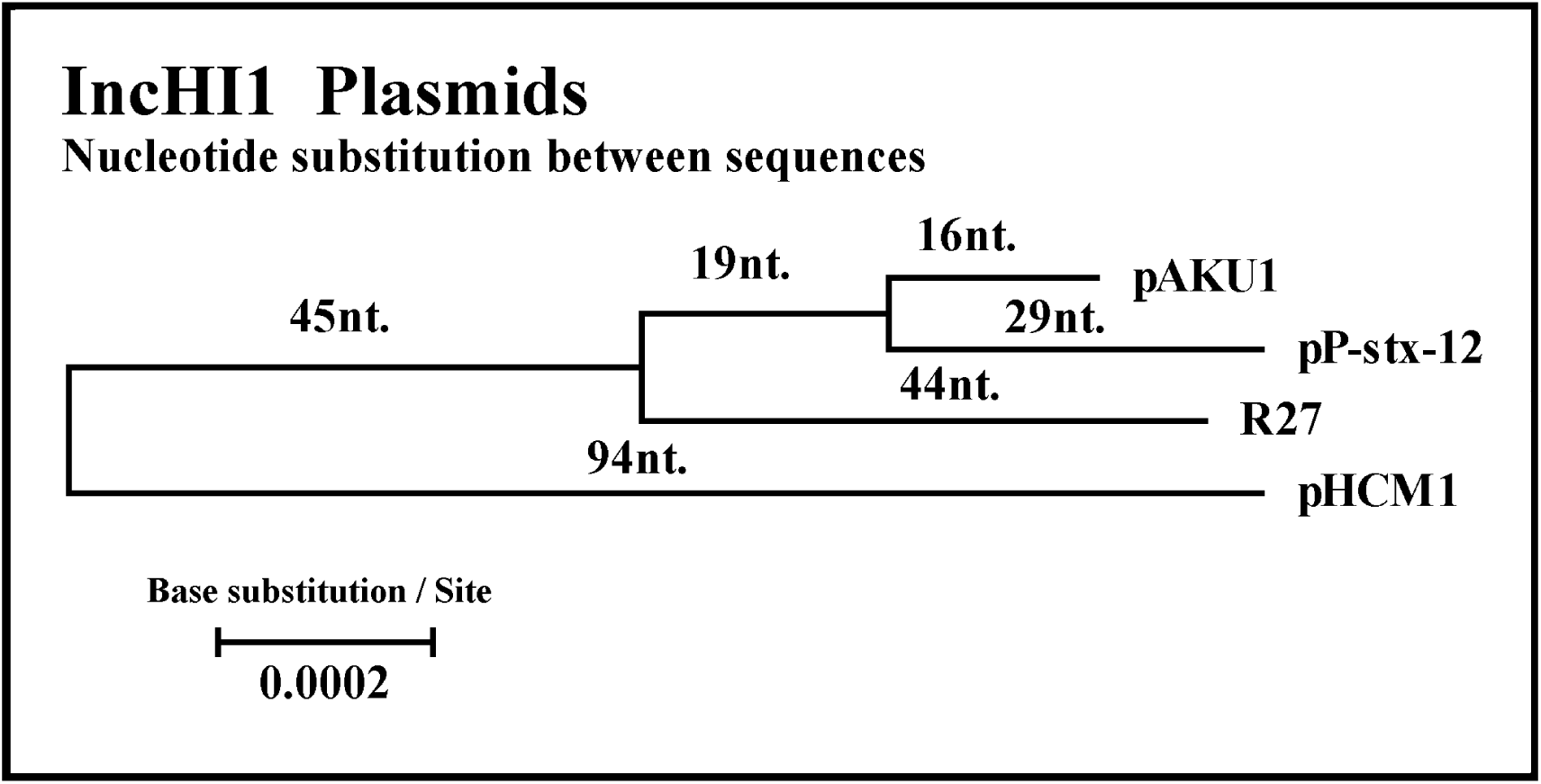
Evolutionary relationships of the taxa and nucleotide substitutions. The phylogenetic tree represents the evolutionary distances, as reflected by nucleotide substitutions, of the extended homologous sequences from the IncHI1 R27-like plasmids (corresponding to the locus R27_p181/192 + R27_p196/207 + R27_p006/21, + R27_p047/73 + R27_p108/122 + R27_p152/168). Base substitutions between the two taxa are the sum of the branches. The substituted nucleotides were determined from the evolutionary distances of the tree; small variations indicate real divergence between the sequences. The final dataset was 85,069 positions (Positions x dS = Nucleotides). Evolutionary analyses used MEGA5. (The tree displayed with BEAST is similar, but pAKU1 and pP-stx-12 show same dS).

The phylogenetical tree showed that pAKU1 and pP-stx-12 homology was related to R27. The pAKU1-like subgroup came, according to the tree, from a 1992 pP-sxt-12-like precursor; analysis by Holt et al. (2011) also found divergence with most recent common ancestor circa 1992 (but R27 was located before both subgroups). In this model, the representative plasmids of both types, pAKU1-like and pHCM1-like, were clearly separated and they were connected by an immediate precursor. Determination of the divergence time (using dS instead of nucleotide substitutions) indicates that the independent evolution of pHCM1 was 48 years (relative to 1960, one year before of the isolation of R27 plasmids); equivalent phylogenetical tree displayed 60 years in the 2011 Holt et al. analysis. Divergence of the others was 23 years later.

Similarly, confirmation of the two types of IncHI1 plasmids was demonstrated by Cain and Hall (2013) based on the presence or absence of variable regions in the DNA. BLASTn analysis of all plasmids confirms Type 2 for pHCM1, pECO111, pMAK1, pEQ2, pF8475, p109/9 and pB71. Type 1 (without BLASTn alignment to variable regions), was confirmed for R27, pP-stx-12, pKU1 and homologues. The pNDM-CIT-like plasmids can be considered a Type 2 lineage since they share segments, but all of the segments have lower identities (from 70% to 96%) and, in some cases, they have short lengths. These observations suggest, as shown in the previous tree display (FIGURE 4), that Type 1 originated more recently than Type 2.

Both of the composite transposon lineages (pAKU1-like and pHCM1-like), observed in the genomics studies were consistent with phylogenetic analysis. Nevertheless, the most relevant differences among derivatives were associated with Tn*10* (with or without *tetD* and *IS10R*). The insertion of the composite transposon in a pHCM1-like ancestor could have moved Tn*10* and motivated its partial deletion; insertion of the composite transposon in a different place in the pAKU1-like precursor (perhaps beside the duplicate IS1; SPAP0175) resulted in Type 1.

In summary –as occurs with composite transposon, with acquisition of resistances-, the evolution of the IncHI1 R27-like plasmids can be inferred, based on the previous comments, as FIGURE 5 suggests.

**FIGURE 5.**
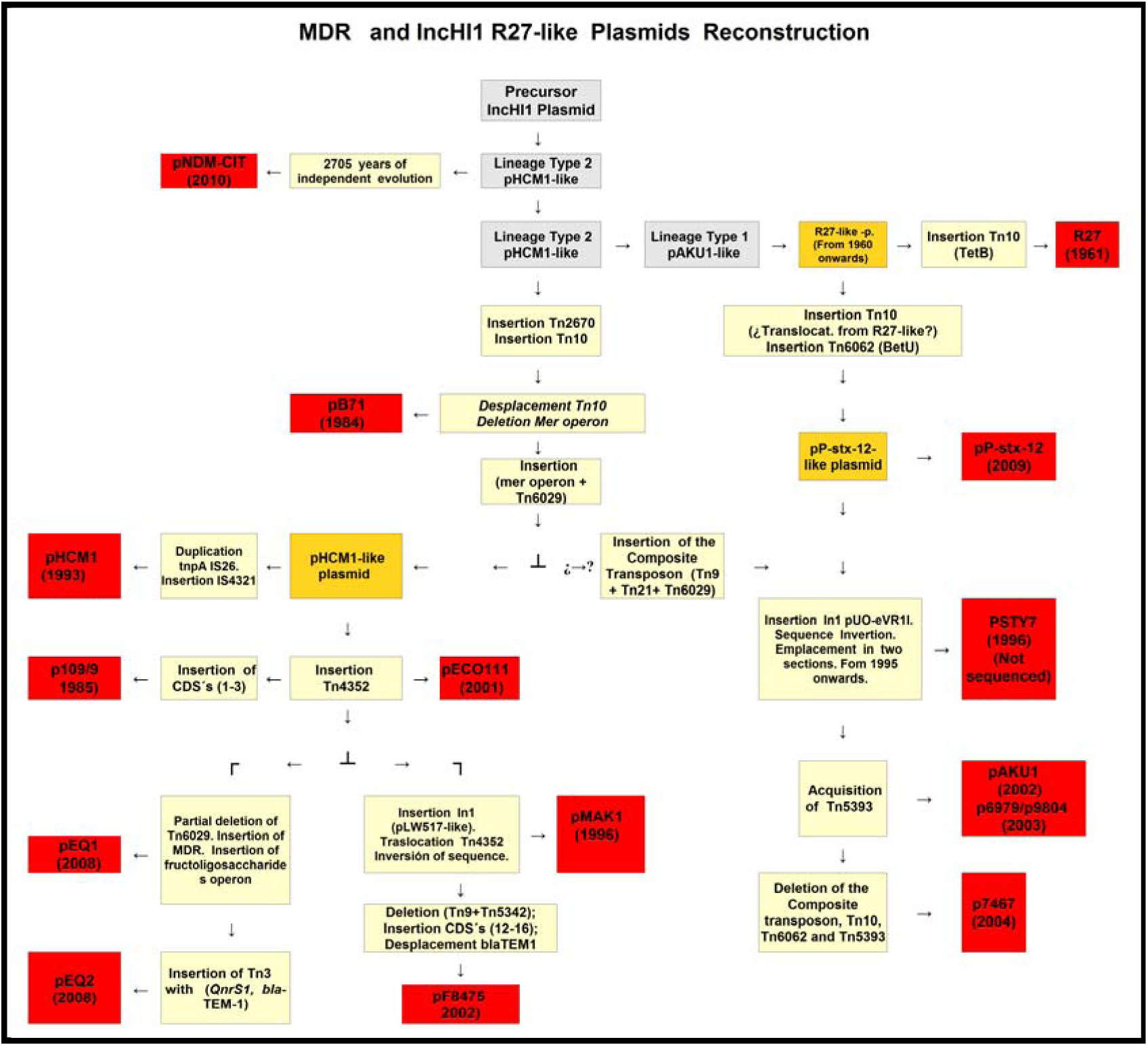
Reconstruction of hypothetical phylogeny of MDR and IncHI1 R27-like plasmids. Plasmids from both subtypes are represented and Type 1 (pAKU1-like) originated more recently than Type 2 (pHCM1-like). Plasmids in red boxes, events in beige boxes. Troubled steps between question marks.

This result does not indicate that each plasmid comes from the predecessor after modification, but the successive acquisition or deletion of transposons, integron cassettes and determinants by the initial composite transposon (as occurs in Tn*4352*, Tn*5393*, Integron pLV517-like, Integron pUO-SeVR1-like, etc.), modulates the development of variations, after which some representative plasmids were isolated and characterized. Reconstruction methodologies are based on the previously noted phylogenetic differences of subtypes and plasmids, as well as MDR loss/acquisition. The events are explained inside the boxes and are described in Supplementary Data 5: “IncHI1 R27-like MDR” and Supplementary Data 2: “Homologous Sequences”.

#### 3.3.3. PHYLOGENETIC RELATIONS AMONG IncHI1 pNDM-CIT-like PLASMIDS

The pNDM-CIT-like plasmids (Supplementary Data 2: “Homologous Sequences”) (excluding the absence of IncHI1 features by simple deletion or by the substitution of segments) showed a few variations from the R27-like subgroup members by the insertion of an MDR module; sixteen CDSs shared with the IncHI2 plasmids remain, as well as the TerZF/TerWY tellurite operon (with 94,5% identity to R478) and the arsenic operon, which is only shared by pNDM-CIT and p8025 (the four different nucleotides with respect to R478, suggests its recent acquisition).

In relation to the IncHI3 plasmids, this clade shares four CDSs; possibly more are shared, but they were not detected with the BLAST options.

In regard to the MDR modules, pNDM-CIT and pF8025 are associated with modules containing different antibiotic determinants, but sharing the initial *IS1/catA1* genes; in other places, they share the phage CP4-like module. The rest of the MDR modules in this subgroup illustrate relations that are consistent with the phylogenetic tree (Supplementary Data 4.-Partial Tree 2). An MDR module with the Tn*1696* Mer operon is shared by pPSP-a3e and pPSP-75c. A long sequence from the genome of the *Enterobacter cloacae* strain colR/S and a second module from Pantotea sp. (and other genomes), both without drug resistances, are common to pKPC_CAV1374, p34983, pPSP-a3e and pPSP-75c. The modules come from different species and horizontal gene transfer can be supposed.

#### 3.3.4. MULTIPLE ANTIBIOTIC RESISTANCE MEDIATED BY IncHI2 PLASMIDS

The IncHI2 MDR modules were not phylogenetically related. The locations and homologies between the heavy metal and antibiotic resistance genes on the IncHI2 plasmids are in Supplementary Data 7: “IncHI HM-MDR”. Each plasmid contains a particular antibiotic resistance gene pattern, encoding different enzymes (Supplementary Data 6: “Antibiotic Determinants”). The heavy metal resistance encoded by the IncHI2 plasmids was related and these results are discussed in the next section (3.3.5).

Other conjugative IncHI2 plasmids were not sequenced but had been identified in strains from many countries. They were partially characterized and, based on their published characteristics, they appeared to belong to the well-characterized lineages R478-like, pCFSAN2050-like or pAPEC-O1-R-like (Cain and Hall 2012a; Cordano and Virgilio 1996; García et al. 2007; García-Fernandez et al. 2007; García-Fernandez and Caratolli 2010; Moreno Switt et al. 2012; Novais et al. 2006; Timme et al. 2013; Zhou et al. 2018).

#### 3.3.5. HEAVY METAL CATION RESISTANCE MEDIATED BY IncHI Plasmids

Some metal cations (Zn^2+^, Cu^2+^, Co^2+^ and Ni^2+^) are essential for living cells and function in enzymatic reactions. Other heavy metal cations (Ag^+^, Cd^2+^, Pb^2+^ and Hg^2+^) are nonessential and toxic. To avoid metal-induced toxicity, microorganisms use a combination of export mechanisms, sequestration and import inhibition at the translational, transcriptional or enzymatic levels.

Mer operon (from Tn*21* or Tn*1696*), is shared by all taxa and clades of the sequenced IncHI plasmids. Except for the subtype R27-like plasmids, the rest of taxa have one or more heavy metal resistance determinants, which are uniformly related to the IncH2 subgroup plasmids. Differences in the IncHI plasmid’s heavy metal resistance operons are in Supplementary Data 8-“HM operons”.

The heavy metal resistance operons were used to determine the relations of the MDR modules in the IncHI2 plasmids, the orientation of the MDR modules, and the phylogenetic events that linked and separated them. The TerZF/TerWY operon was assigned to the extensive core from the common ancestor of IncHI2 subgroup. The Kla operon (also for tellurite resistance), which was partially deleted in some plasmids, displayed complete identity between them (except for pCFSAN2050-1 with 95% identity); the same pattern occurred with theTn*2501*/Tn*2502* arsenical resistance operon. (Tn*2501* is shared by the IncHI3 pKOX-R-like plasmids and by the IncHI3 pNDM-MAR-like plasmid pKpn23412, but is placed inside the MDR module). Other heavy metal resistance determinants that can be found in plasmids are lead (*pbrABCR* genes), nickel (*ncrCR* genes), cation efflux protein (*czcC* gene), copper tolerance (*cutA* gene), silver tolerance (*silA* gene), or magnesium influx (*corA* gene).

##### 3.3.5.1. Triple operon copper resistance/silver resistance/Tn*7*

BLASTn analysis with the triple operon copper/silver/Tn*7* show remarkable results. A complete 32-kb segment was identical in R478, pSH111_227 and pAPEC-O1-R, with nearly 100% identity between them. The differences were so minimal (for an IS4 in APECO1_O1R103, 14-nucleotide difference in pSH111 and 10-nucleotide difference in pAPEC-O1-R with respect toR478) that a common ancestor seems plausible, even though R474 is located in a different clade in the phylogenetic tree (Supplementary Data 4.- Partial Tree 3). An exact insertion in the same position and with the same homology is difficult to explain, especially because R478 was isolated in Boston in 1969 from a clinical isolate of *Serratia marcescens* (Madeiros and OBrien, 1969), pAPEC-O1-R was isolated in Arkansas in the early 1990s from an avian pathogenic *Escherichia coli* isolate (Johnson et al. 2006), and pSH111_227 was isolated from *Salmonella enteric* in a cow from Ohio in 2001 (Han et al., 2012). Although the acquisition of the triple operon segment might not have been random in the three plasmids, its perfect homology did not evolve in parallel with the plasmids (in the Tra1+Tra2 studies, homologues have 98.3% identity; up to a 250 nucleotide difference in a total of 10,132 positions in the final dataset).In the absence of a better explanations (as exiguous mutation rate), the improbable convergent acquisition by horizontal gene transfer, or the recombination between shared core regions flanking the triple operon with compatible plasmid should be considered.

The other nine plasmids of the IncHI2 subgroup without this triple module have some CDSs in the same location. Two of these CDSs, located in the middle of the sequence, correspond to the two first ORFs of the Cop operon; besides *copE1*, these plasmids gained the NcrC and YohL determinants for nickel resistance (Rodrigue et al. 2005). This result suggests that a common ancestor suffered a deletion of the triple operon and the posterior insertion of the new sequences. However, BLASTn analysis did not confirm this result since the identities of *copE1-copS*, with respect to R478, are approximately 95%.

pCFSAN2050-2, which is displayed as a clade in the phylogenetic tree (Supplementary Data 4.- Partial Tree 3), has *copE1-copS* and the surrounding genes and, after a MDR module, it has the last gene of the silver resistance operon and the Tn*7*. This sequence is shared with 99,9% identity in the complete genomes of different bacteria (as studied by carbapenemase production in Conlan S et al., 2014 and Sheppard AE et al., 2016). The high similarity of these sequences and the low homology to the triple operon (96% identity with respect to the same segment in R478), point to independent acquisition and not a common ancestor with R478 or its homologues. These identities, which are similar in the *copE1-copS* determinants, suggest that there was a precursor with an equivalent of the triple operon module. In addition, all of the plasmids with *copE1-copS* determinants were isolated since 2001, when the last plasmid with the complete triple operon was isolated. Supporting this supposition, in the plasmid pP10164-2, studied by Sun et al (2016), beside *rcnR/pcoE1/pcoS* genes (99,9% of identities with homologous in pCFSAN2050-2), is emplaced the complete Silver operon (94% of identities with R478). In plasmid pT5282-mphA, studied by Liang et al (2017), remains only *silR* gene (100% of identities with pP10164-2).

In conclusion, the complete triple operon and the segment with the *rcnR/pcoE1/pcoS* genes (identified in the plasmids of the same clade) could have different origins, but the same location.

##### 3.3.5.2. Mercury resistance operon

A Mer operon can be detected on 32 plasmids belonging to the IncHI1, IncHI2 and IncHI3 subgroups.

The Tn*1696* Mer operon was not distinctive to the IncHI2 subgroup since it is absent in some plasmids and has different location in others, suggesting a recent acquisition and not a parallel evolution from an R478-like ancestor. The plasmids pEC-IMP/pEC-IMPQ, pCAV1151, pKPC-272, pENT-8a4 and pMRVIM0813 also had a truncate transposon with Mer and Lead determinants that was the derivative of a transposon related to Tn*1696* in R478. This transposon was surrounded by 5-bp duplication, indicating that it was acquired via transposition from a different origin (Cain et al. 2010; Cain and Hall 2012a).The Tn*1696* Mer operon in pCFSAN2050-1was emplaced in different location; the plasmids pSSTm-A54650 and pKST313 had a Tn*21* Mer operon inside the MDR modules as well.

The IncHI1 R27-like clade shared the Tn*21* Mer operon inside the composite transposon. However, pHCM1 also had a Tn*1696* Mer operon placed in an equivalent position to the IncHI2 plasmids, but in the opposite direction and from a different origin. In the pNDM-CIT-like clade, the Tn*1696* Mer operon was present inside a similar MDR module in pPSP-a3e and pPSP-75c; the plasmid pF8025 had a Tn*21* Mer operon.

The IncHI3 plasmids shared a module with the Mer operon. Curiously, in the pNDM-MAR-like clade, two of the plasmids had a Tn*21* Mer operon, three had the Tn*1696* Mer type, and one (pENVA) had a different Mer operon.

##### 3.3.5.3. Tellurite resistance operon (TerZF/WY) phylogeny

To clarify the phylogenetic relationship of the Ter operon, molecular clock analysis was used with the tellurite resistance operon genes. The results of the BEAST phylogenetic tree are in FIGURE 6.

**FIGURE 6.**
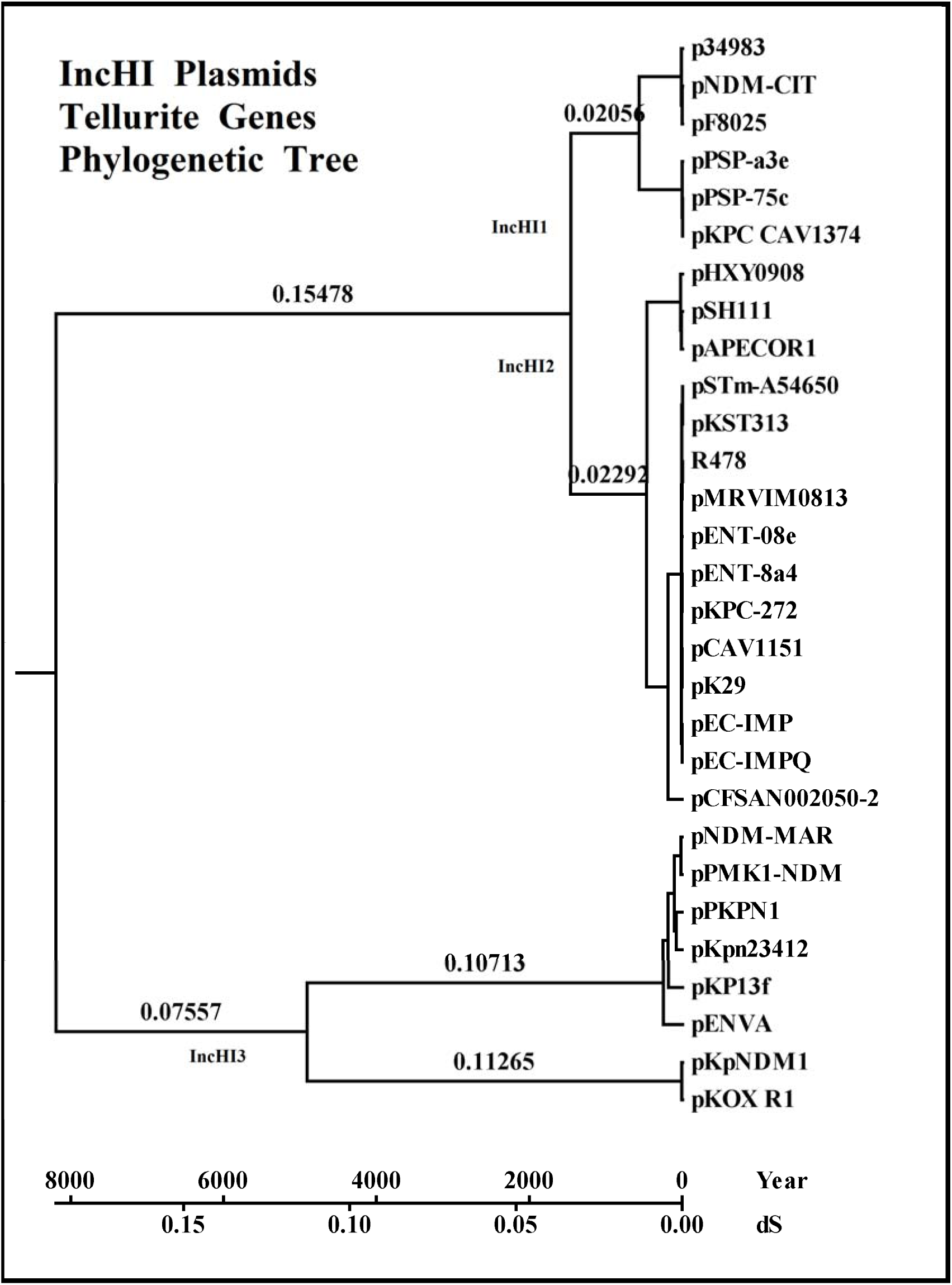
Evolutionary relationships of taxa related to the tellurite resistance operon of IncHI plasmids. Correspondence is between the number of bases substituted per site and the years of divergence, with 7,298 positions in the final dataset. Evolutionary analyses used the BEAST package. The tree was drawn with MEGA5 software.

The taxa and clades displayed in the tellurite resistance gene tree (FIGURE 6) are identical to those in the Tra gene tree (FIGURE 1), but with different divergence times. The estimated complete divergence from a common ancestor corresponds to 17,509 years in Tra genes and 8,150 years in tellurite genes. Theoretically, this difference suggests that the Ter operon was acquired after the divergence in the taxon but, with the trees being so similar, the most feasible hypothesis is that the operon was joined to the IncHI group before this divergence (including the IncHI1 R27-like plasmids, which lost the operon, possibly due to their divergence with the pNDM-CIT-like plasmids). Consequently, it can be assumed that clock rate (μ= mutations per nucleotide per year), of the tellurite resistance operon is different from that of the Tra genes.

The partial tree of each taxon (Supplementary Data 9-“Tellurite Genes Sub-Trees”) does not clarify divergences. The IncHI2 plasmid tellurite resistance genes tree (Supplementary Data 9-Partial Tree 2), with a divergence time of 467 years, has two times the display of transfer genes (226 years), but divergence is caused mainly by clade pSH111/pAPECOR1/pHXY0908.

Two related clades are displayed in the IncHI1 pNDM-CIT-like tellurite resistance tree (Supplementary Data 9-Partial Tree 1). The divergence time of the tellurite resistance genes was 570 years, whereas it was 13,5 years in the Tra genes; both trees coincide in the clade with plasmids pPSP-75c, pPSP-a3e and pKPV ACV1374, but not in the other one. Until a better explanation can be found, this result suggested an unusual SNP variation in the common ancestor of the clade or the improbable acquisition of a novel complete TerZF/TerWY operon (improbable because the adjacent haloacid dehalogenase, HAD, which is also shared by the IncHI3 plasmids, still remains).

The IncHI3 taxon (Supplementary Data 9-Partial Tree 3) of the tellurite resistance genes tree is similar with respect to the Tra genes tree for both clades (4,912 years of divergence for Ter genes and 4,512 years of divergence for Tra genes). Differences with respect to the clades are from patents. For the IncHI3 pKOX-R1-like plasmids (which both have deletion of the TerWY operon), the 207 years of divergence in the Tra genes was reduced to 9 years in the Ter genes. With the IncHI3 pNDM-MAR-like clade, the transfer genes display a tree with 627 years of divergence and tellurite resistance genes display 292 years of divergence.

In conclusion, the tellurite resistance operon, as component of the IncHI core, has a tree topology that mirrors the overall plasmid-backbone tree, but it has some differences in its mutation rate and divergences in its sub-trees and branches.

#### 3.3.6. Phylogenetic relations among IncHI2 plasmids

The suggested IncHI2 plasmid evolutive reconstruction is shown in FIGURE 7.

**FIGURE 7.**
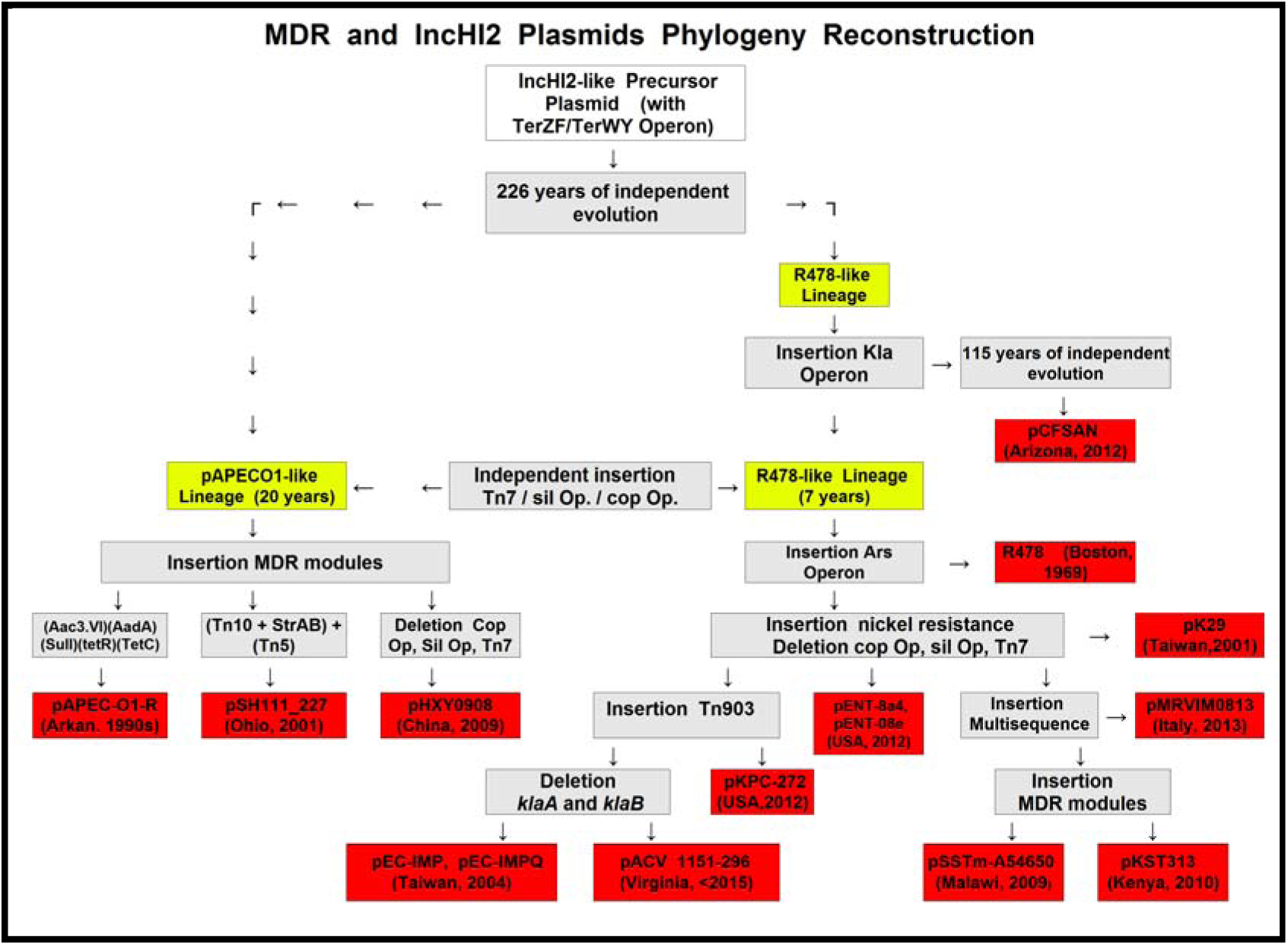
Heavy metal operons and IncHI2 phylogeny reconstruction. Age was calculated from the IncHI2 Tra genes (Supplementary Data 4-Partial Tree 3), and refers to the years of divergence. Evolutionary relationships among the lineages is represented. Plasmids appear in red boxes, events are in grey boxes.

The relationship between the IncHI2 plasmids, according to Tra genes homology, core similarity and MDR modules, is thus established. Nevertheless, plasmid evolution progresses through recombination with the MDR determinants, not through the accumulation of SNPs in a short time within a close ecological niche (Fricke et al. 2009).

Reconstruction methodologies are based on phylogenetic tree results (Supplementary Data 4-Partial Tree 3), heavy metal operon differences (Supplementary Data 8- “HM operons”) or MDR module variations (Supplementary Data 7: “IncHI HM-MDR”). The events indicated two main clades. The pAPECO1-like lineage (with 226 years of independent evolution with respect to the R478-like lineage) showed three plasmids with very different MDR modules, variations in the HM operons, and core differences that did not justify the displayed branch of the tree. This lineage has sustained an independent evolution from R478-like lineage, after acquisition, by R478 ancestor, of KlaABC. pCFSAN2050 has sustained an independent evolution from the R478-like lineage, according with the tree, 115 years ago. R478 plasmid appeared in 1969, and the next isolated plasmid in this linage was pK29 in 2001, which does not have the triple operon, but *copE1 and copS* remain. The insertion (as Tn*903*) or deletion (as KlaAB) of significant sequences clarified the reconstruction of the lineage evolution and the relations between plasmids.

The three IncHI2 plasmids studied by Liang et al (2017) and by Sun et al (2016), carry in same position the heavy metal resistances (Mer operon, Ter operons, Kla operon, Ars operons, and *RcnR/PcoE1*/*PcoS* genes, but not the Pbr operon), as well as an array of antibiotic resistance genes located in MDR modules and associate with mobile elements. Each plasmid has peculiarities.

In the plasmid pT5282-mphA (GenBank: KY270852), remains silR (94% identities with R478), insides the module adjacent to RcnR/PcoE1/PcoS determinants, but without antibiotic resistance genes. The plasmid contained four accessory resistance regions.

In the plasmid p112298-catA (GenBank: KY270851), the regions with RcnR/PcoE1/PcoS and Tn1696-Mer operon are absents, but remains the MDR module adjacent to Kla operon. Most of the segment is shared by pEC-IMPQ plasmid but located in different sections and with disruptions.

The plasmid pP10164-2 (GenBank: KX710093), is almost identical in core sequence to plasmid pKST313, but different in accessory regions. The only MDR module, with 98 Kb, is made up by the union of the two MDR initial sections (because the DNA between both was deleted); one section becomes from the *RcnR/PcoE1*/*PcoS* MDR region (pKSt313, *CDSs* 324-321), and the second one from the MDR region adjacent to helicase gene *yjcD* (pKSt313, *CDS* 257).

In sum, the MDR modules from new studied IncHI2 plasmids become from similar origin with different insertion sequences associated with integrons, transposons and mobile elements, harboring genes conferring resistance to multiple classes of antibiotics.

#### 3.3.7. IncHI3 PLASMIDS

The IncHI3 plasmids came from the ancestral lineage of the IncHI group, with notable relevant characteristics. The homology of their shared features with the other IncHI plasmids showed a clear relationship; this relationship was confirmed by protein alignment because the nucleotides did not display significant homology with BLASTn (Supplementary Data 2: “Homologous Sequences”; pNDM-MAR plasmid, CDS homology). This correspondence pointed to a phylogenetic divergence, supported by distance from a common ancestor that was a consequence of evolutionary differentiation. Phylogenetic distance among clades is reflected in core differences. Two to four MDR modules are present in each plasmid, and some of the MDR modules are shared by a few plasmids. Tn*10* (*tetB*) is shared by the IncHI3 pKOX-R1-like clade.

The Mer module, positioned below the TerWY operon, is an example of the MDR module evolution and HGT. Previous to the Mer operon, a sequence with the *yadA* gene is shared by the IncHI3 pNDM-MAR-like plasmids. pKP13f, isolated in 2009, has the Tn*21* Mer operon. pKpn23412, isolated in 2011, is derived from a common ancestor, with the same transposases flanking the operon, but with the substitution of a sequence (Tn*2502* and Tn*9* between them) for the *yadA* gene. As well, after the Mer operon, there is the addition of Tn*6026* and other determinants in pKpn23412. A second derivative from pKP13f-like is pNDM-MAR, isolated in 2010; it has a *yadA* gene, but the Tn*1696* Mer operon replaced the Tn21 Mer operon and, after a novel additional sequence, it includes the initial 4800 bp of Tn*1696*. pNDM-MAR-like is the beginning of two new lines of plasmids. The first line contains pPMK1-NDM, isolated in 2011. It has a novel insertion of multi-sequences adjacent to the previous sequence. pPKPN1, isolated in 2013, is a derivative of this plasmid and has a partition that separates the module into two sections: an inversion of one sequence with its adjacent genes and the addition of determinants. The second line corresponds to plasmid pENVA, isolated in 2012, which includes the initial 4800 bp of Tn*1696* in a module with newly acquired determinants and, on the other side of an inversion, the *YadA* sequence with substitution of the Tn*1696* Mer operon and its adjacent sequences with a truncate Mer operon (99,9% identity to *A*.*baumannii* 1656-2).The reconstruction of phylogeny is shown in FIGURE 8.

**FIGURE 8.**
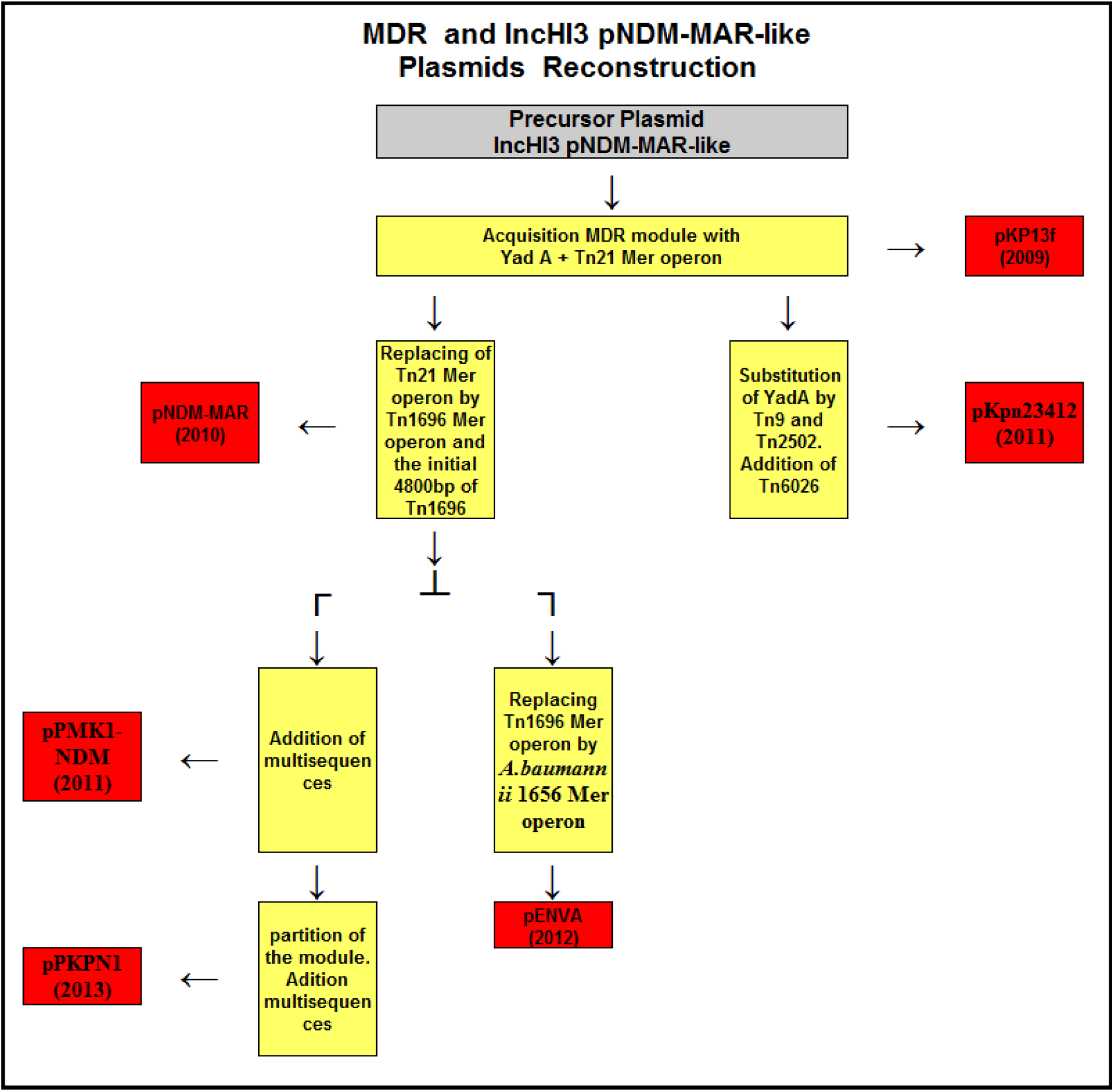
MDR and IncHI3 pNDM-MAR-like plasmids reconstruction of hypothetical phylogeny. The diagram is based on the*YadA* gene and Mer operon from the MDR modules. Plasmids appear in red boxes, events in yellow boxes.

The two IncHI3 pKOX-R1-like plasmids shared (in same location, besides the truncated TerWY operon) an equivalent MDR module with the Tn*1696* Mer operon (truncated in pKpNDM1 by the insertion of sequences) and the initial 4800 bp of Tn*1696* in another module. Interestingly, Tn2501 (Ars operon) is present in both plasmids of this clade, inside the same MDR module, and Tn*10* (TetR) is present, all alone, in the core.

The plasmid pYNKP001-dfrA, recently studied by Liang et al (2017), with a core similar to the IncHI3 pKOX-like plasmids, have the same truncated TerWY operon and only the MDR module adjacent to *terW*. As pYNKP001-dfrA plasmid have not partial inversion of the core (as occurs with pKOX plasmids), is more similar to pKpNDM1plasmid. The antibiotic resistance region star with the same CDSs and the complete Tn*1696* mer operon of pKOX plasmid, and continues with a different array of sequences inserted.

### 3.4. H-Group PLASMID BACKBONE

The MPF genes of all sequenced IncHI plasmids resembled, in terms of genetic organization and sequence identity, those encoded on the F plasmid (Supplementary Data 3: “Backbone”). Other conjugative elements also shared the same MPF determinants (Gilmour et al. 2004; Lawley et al. 2003a).Within the H plasmids, the coupling protein TraG and the relaxosomal proteins TraI and TraJ constituted a novel phylogenetic group titled MOB_H_; this group had multiple conserved genetic elements (Garcillán-Barcia et al. 2009; Guilmour et al. 2004) and could be transferred as a mobilization module. These observations suggest that the MPF and MOB determinants of IncH plasmids evolved from different lineages (Gilmour et al. 2004). The highly related H-group elements SXT (ICE), R391(IncJ/ICE), Rts1(IncT), pCAR1(IncP7), pSN254 (IncA/C) and RA1 (IncA/C) encoded both an F-like transfer system and MOB_H_ mobilization system, but did not share the core determinants of the IncHI plasmids. Molecular clock analysis was used to determine phylogenetical relationships between tra genes.

To determine phylogenetic relationships among these plasmids and the IncHI group, the nucleotide sequences from 15 conjugative transfer system genes were downloaded from the European Nucleotide Archive (http://www.ebi.ac.uk/ena/) and the concatenated sequence of each was used to determine distance estimation and divergence time. The results of the Bayesian evolutionary analysis with BEAST are shown in FIGURE 9.

**FIGURE 9.**
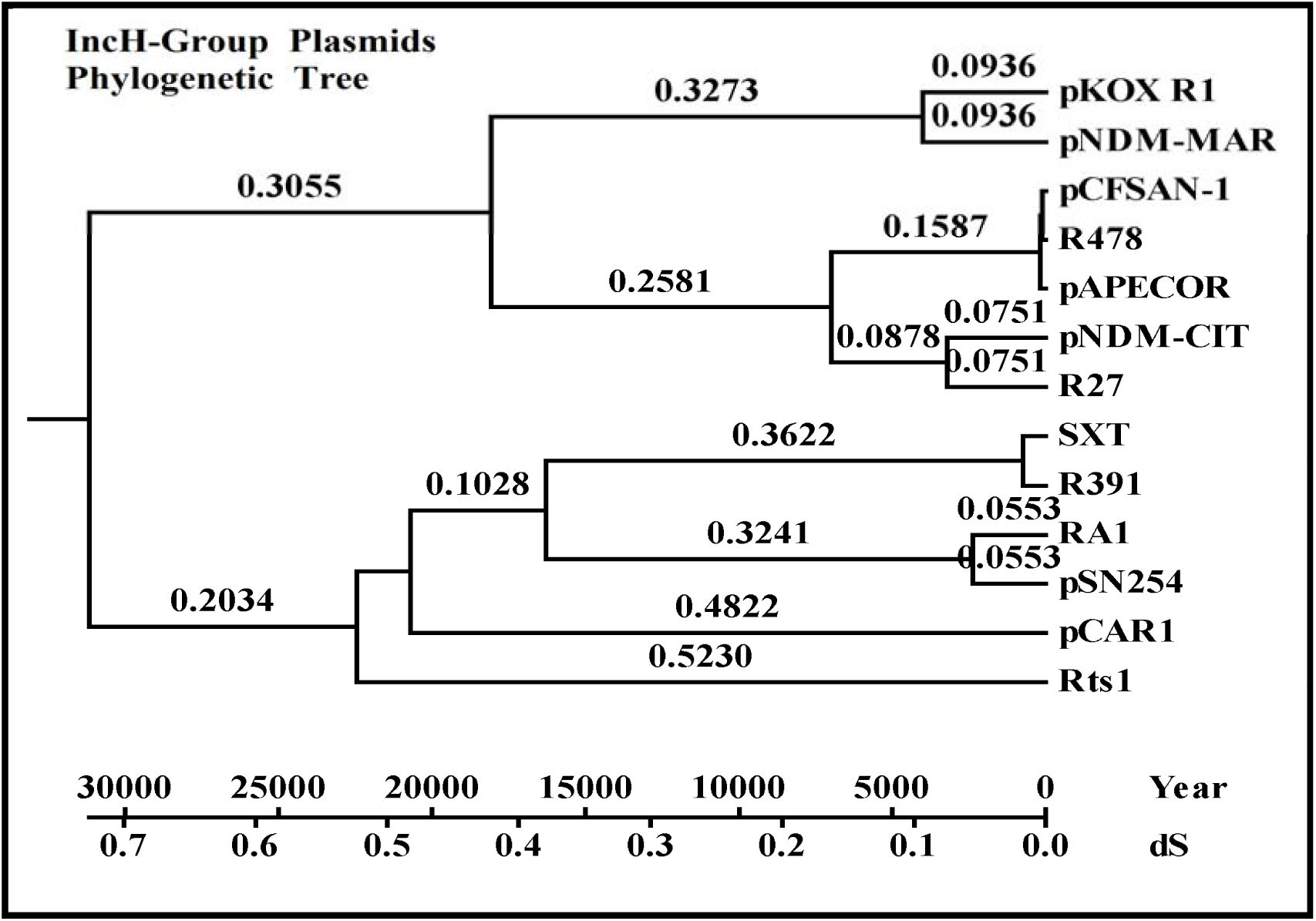
Evolutionary relationships of H-Group taxa inferred using the minimum evolution method. The tree shows the correspondence between synonymous substitution and the years of divergence. Branch lengths refer to number of synonymous substitutions per synonymous site for 17,355 positions in the final dataset. Evolutionary analyses used BEAST. The trees were drawn with MEGA5.

The divergence time of the tested genes in the H-group plasmids from a common ancestor was estimated at 31,617 years. The H-group related elements are as interrelated as the IncHI plasmids, so a common ancestor can be accepted for tra genes. The accuracy of the results for this tree could be extended to the previous tree (FIGURE 1). IncHI plasmids (IncHI1+IncHI2+IncHI3) display a divergence of 18,320 years (17,509 years in FIGURE 1); IncHI1+IncHI2 plasmids display a divergence of 7,090 years (8,176 years in FIGURE 1). pNDM-CIT and R27 display a divergence of 3,269 (2,705 years in FIGURE 1).

## 4. CONCLUSIONS

The results presented here provide insights into IncHI plasmid tra genes phylogeny. A relationship among the transfer regions of the H-group plasmids was clear, with divergence from a common ancestor nearly 31,617 years ago. The IncHI plasmid subgroups IncHI1 and IncHI2 shared a common ancestor around 8,176 years ago. The homology of the core sequences in the IncHI1 subgroup confirmed these close relationships and the remarkable divergence of the pNDM-CIT-like lineage plasmids 2,705 years ago. The IncHI2 subgroup had proximal homology in all plasmids. Analysis of the IncHI2 Tra genes by molecular clock test clearly showed that R478 and nine other plasmids belong to the same lineage, whereas pAPEC-O1-R, pSH111_227 and pHXT0908 belong to a different lineage from 226 years ago; pCFSAN2050 came from an ancestral R478-like plasmid 155 years ago. The IncHI3 plasmids (pNDM-MAR-like/pKOX-R1-like) were different; only 105 core determinants were shared with rest of the IncHI plasmids, which shared 130 determinants. However, five CDSs from IncHI3 were shared exclusively by the IncHI1 subgroup and four were shared by the IncHI2 subgroup. BLASTn identities from common IncHI3 with homologous IncHI1/IncHI2 determinants varied from 25% to 75%. This result suggested that IncHI3 came from a common ancestor with the other IncHI plasmids 17,509 years ago.

Most IncHI1 R27-like subgroup plasmids shared a controversial composite transposon. Two transposon lineages were proposed that differ in the presence or absence of the resolvase TnpR from Tn*21*. However, the adjacent multidrug-resistance determinants came from a characterized integron and were confirmed by a derivative composite transposon with complete *tnpR* that was identified in an *S*.*typhi* genomic island. However, the old plasmid pB71 (isolated in 1984) had *IS, catA1*, and the initial 8,000 bp of Tn*21* (including *tnpR*) as a proto-transposon, without the rest of the transposons and Mer determinants. pB71-like could be a feasible precursor for the group without *tnpR* since a truncate Tn*10* is located beside IS1. Tn*10* is the most feasible module to differentiate both clusters, since it has different locations and is not truncate in plasmids with *tnpR*. The evolutionary relationship of a common core segment of 85K bp from representative IncHI1 R27-like plasmids (FIGURE 4) showed that nucleotide substitutions between sequences were duplicated in plasmids with truncate Tn*10* and without *tnpR* in the composite transposon; this result indicates that plasmids with *tnpR*_*Tn21*_ came later.

The IncHI1 plasmids pNDM-CIT-like represented a different IncHI1 lineage with respect to the R27-like subgroup; they share different MDR modules and some CDSs related to the IncHI2 plasmids, such as the RelBE module, arsenic operon and the TerZF/TerWY (in different location).These results suggested that the pNDM-CIT lineage came from a different ecological niche than the R27-like plasmids. In addition to these peculiarities, pNDM-CIT-like plasmids have an IncHI1 core with characteristic variable regions for the Type2 pHCM1-like plasmids (without *tnpR*), which corroborates that Type1 (R27/pAKU1-like) originated more recently than Type2.

The IncHI2 MDR modules did not have phylogenetic relationships, but its core heavy metal resistance genes and operons were related. The most relevant, the TerZF/TerWY operon, is shared by all IncHI plasmids (except IncHI1 R27-like plasmids), with different identities in taxa and clades according to phylogenetic Tra gene evolution. Acquisition by a common ancestor and parallel evolution (with a different mutation rate than Tra genes) seems to be the most likely interpretation, in spite of the fact that some sub-tree clades and branches display unexpected divergence. Even though, this unexpected divergence does not support the unlikely case of independent acquisition after cluster segregation.

The extended segment with nickel resistance, the copper resistance operon, the silver resistance operon and Tn*7* showed unusual results. The complete sequence of this triple operon was shared by plasmids R478, pSH111_227 and pAPEC-O1-R, although the transposase IS4 was inserted in APECO1_O1R103. Parent strains from each plasmid were isolated in a 32-year period and displayed only ten different nucleotides in a 32-kb sequence. No rational explanation could be found.

In remaining IncHI2 plasmids, only two copper determinants (with 96% identity to same determinant in R478) were detected. The defective homology and the addition of nickel resistance suggest that different origins is more feasible explanation; this explanation is supported by the pCFSAN2050-2 plasmid, which also shares *copE1-copS* and the surrounding genes and has, after an inserted MDR module, the last gene of the silver resistance operon and Tn*7*, with identities of 96% to R478,which point to an equivalent triple operon.

Finally, the IncHI3 plasmids had lower homology in the Tra regions than the IncnHI1 and IncHI2 subgroups, and this result clarified the IncHI phylogeny since it seemed to come from a common ancestor of the group before the differentiation of lineages. The core ORFs linked the IncHI3 plasmids to the rest of the IncHI plasmids, but their poor homology and limited number of common determinants had sufficient distances to prevent their incorporation into the phylogeny of the IncHI1 and IncHI2 subgroups; instead, they were included as a subgroup IncHI3.

In summary, sequence analysis revealed that each of the IncHI plasmids retained a conserved backbone sequence and extensive homology in core regions. Differences were primarily within the accessory segments, which appeared to evolve via horizontal gene transfer events. IncHI plasmid analysis suggested an evolutionary model in which each subgroup derived from a common ancestor through a process of stepwise modification events in the core determinants and, more recently, horizontally acquired resistance gene arrays on plasmids. Although the evolution of plasmids by the accumulation of point mutations is a natural consequence of time, and the acquisition of resistance elements depends on the environment, both properties contribute to variations from common ancestors.

## Supporting information

plasmids

Tra genes

Genbank datta

Homologous sequences

Backbone

Tra genes sub-tree

IncHI1 R27 MDR

Antibiotic determinants

IncHI HM-MDR

HM Operons

Tellurite genes sub treee

## GLOSSARY

CDS: Coding DNA Sequence (Portion of a gene’s DNA or RNA, composed of exons, that codes for protein. While identification of open reading frames within a DNA sequence is straightforward, identifying coding sequences is not, because the cell translates only a subset of all open reading frames to proteins).
dS: Distance estimation (Synonymous substitution rate; numbers of synonymous substitutions per synonymous site).
HGT: Horizontal Gene Transfer. (The movement of genetic material between bacteria other than vertical gene transfer, in which information travels through the generations as the cell divides).
ICE: Integrating Conjugative Elements are mobile genetic elements that can transfer from the chromosome of a host to the chromosome of a new host through the process of excision, conjugation, and integration.
MDR: Multi Drug Resistance. (A condition enabling a disease-causing organism to resist distinct drugs or chemicals of a wide variety of structure and function targeted at eradicating the organism).
MOB: Mobilization machinery (which encodes a complete set of transfer genes, the cytoplasmic nucleoprotein relaxosome complex)
MPF: Mating-Pair Formation. (A set of proteins that elaborate the transport channel as well as a pilus that allows the attachment to the recipient cell, and thereby the translocation of the relaxase-DNA complex.
ORF: Open Reading Frame. (Part of a gene that encodes a protein)
SNP: Single-Nucleotide Polymorphism. DNA sequence variation occurring when a single nucleotide in the genome (or other shared sequence) differs between members of a biological species or paired chromosomes in an individual.

