## Supplementary material for "Comparative genomics and phylogeny of sequenced IncHI plasmids": Tra genes sub-tree

**Tra GENES SUB-TREES**

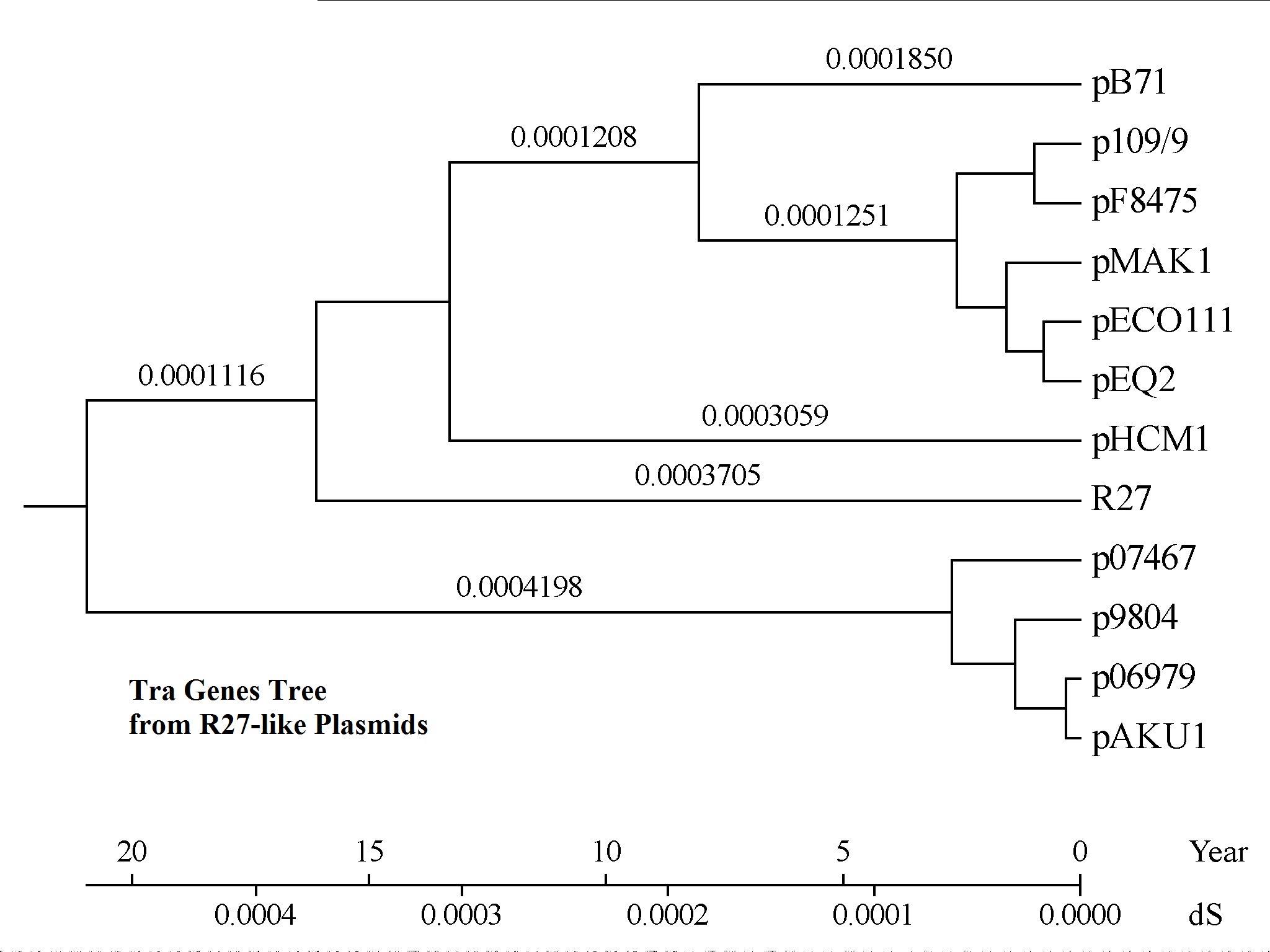

**Tra Genes Sub-Tree 1a**.- IncHI1 R27-like plasmids Tra genes tree.

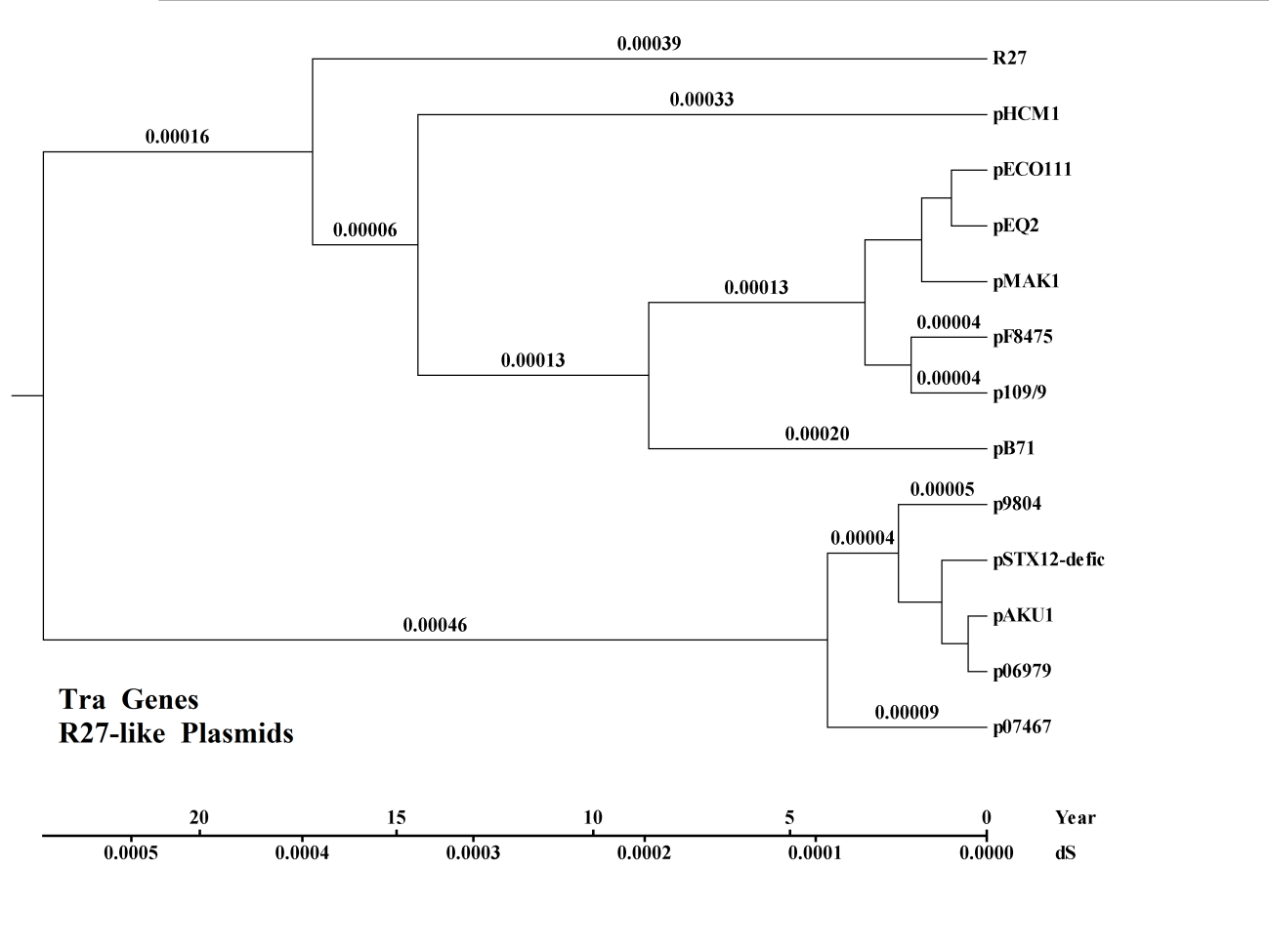

**Tra Genes Sub-Tree 1b**.- IncHI1 R27-like plasmids from the expanded FIGURE 1.

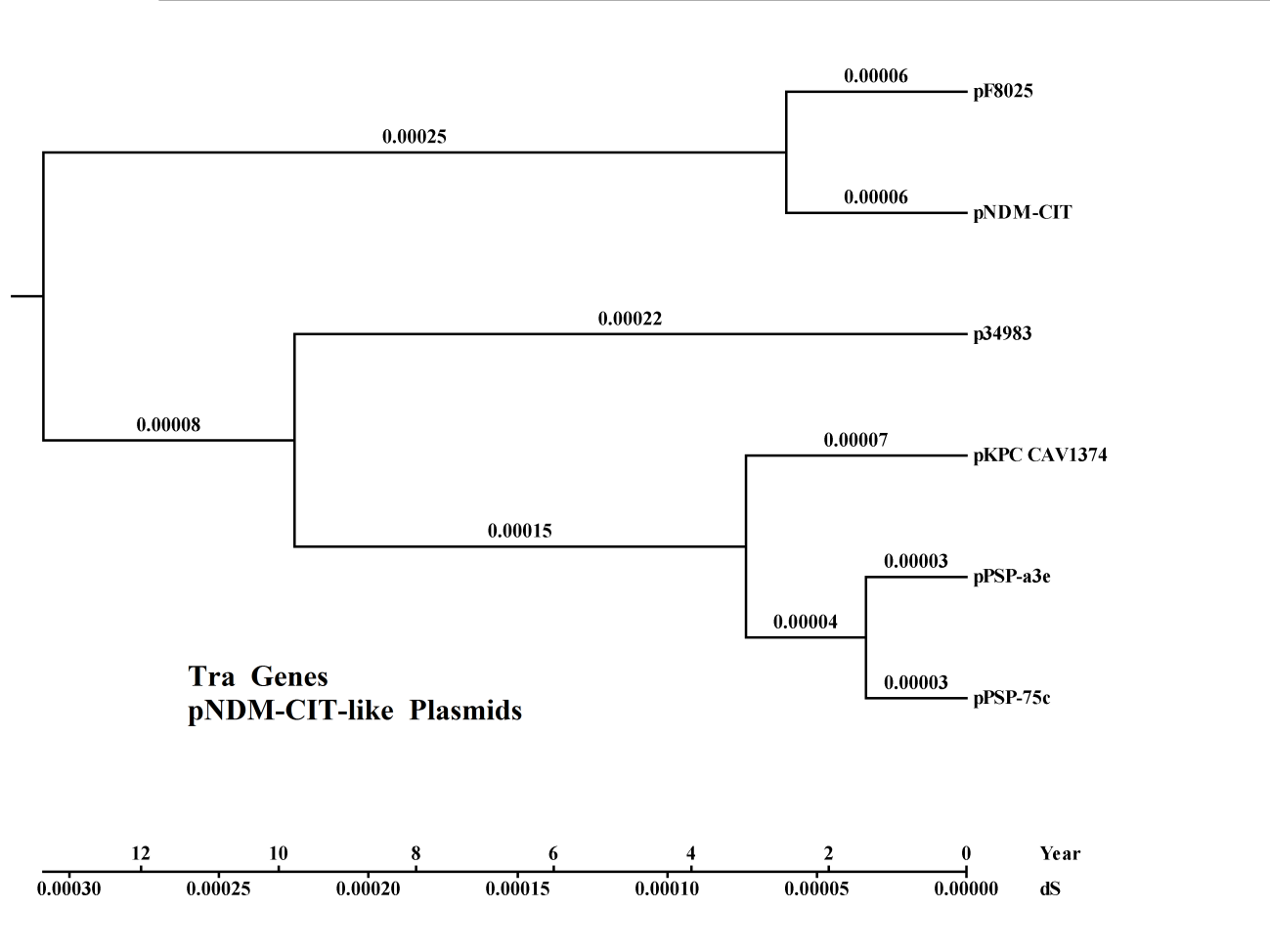

**Tra Genes Sub-Tree 2**.- IncHI1 pNDM-CIT-like plasmids from the expanded FIGURE 1.

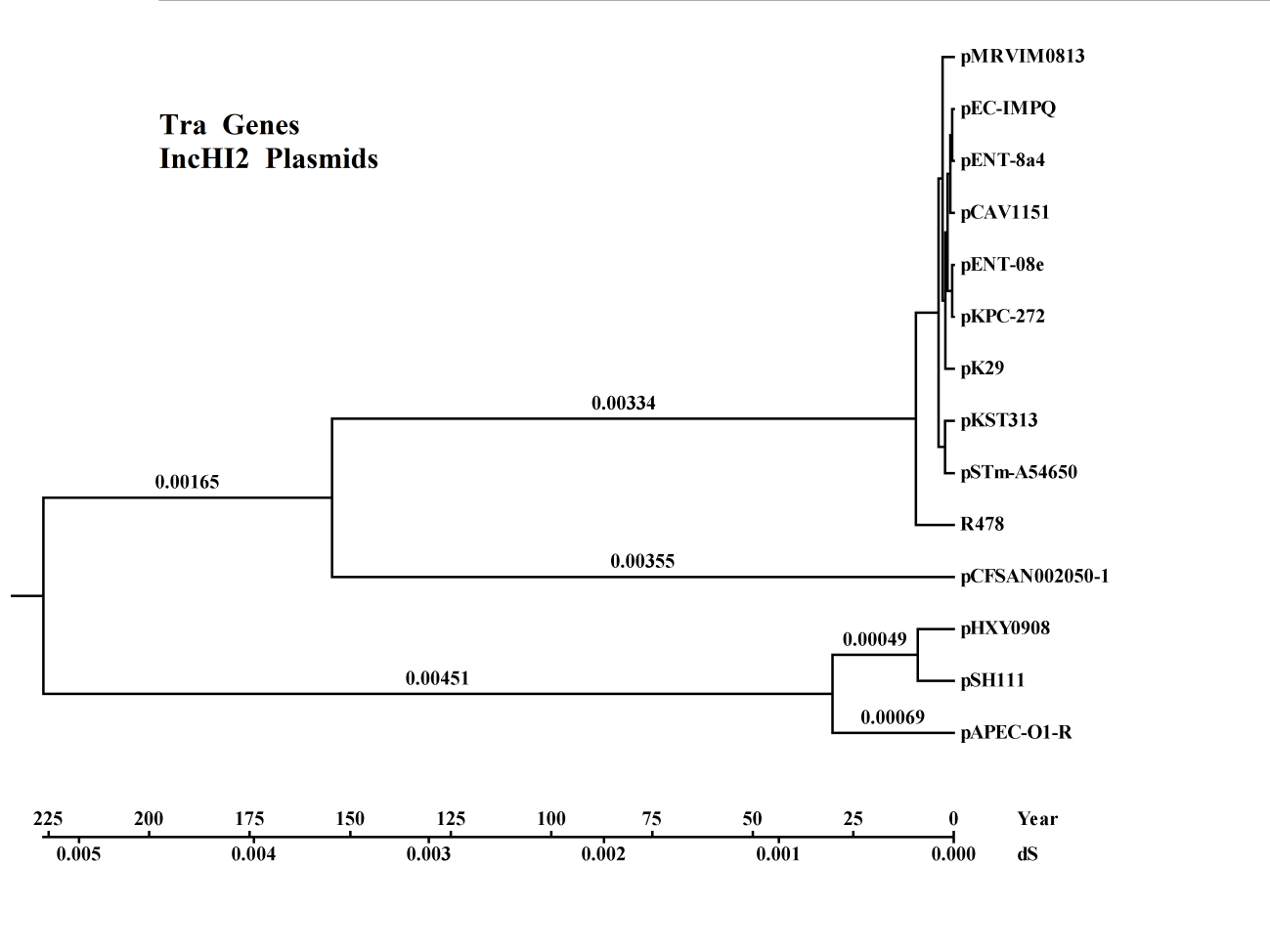

**Tra Genes Sub-Tree 3**.- IncHI2 plasmids from the expanded FIGURE 1.

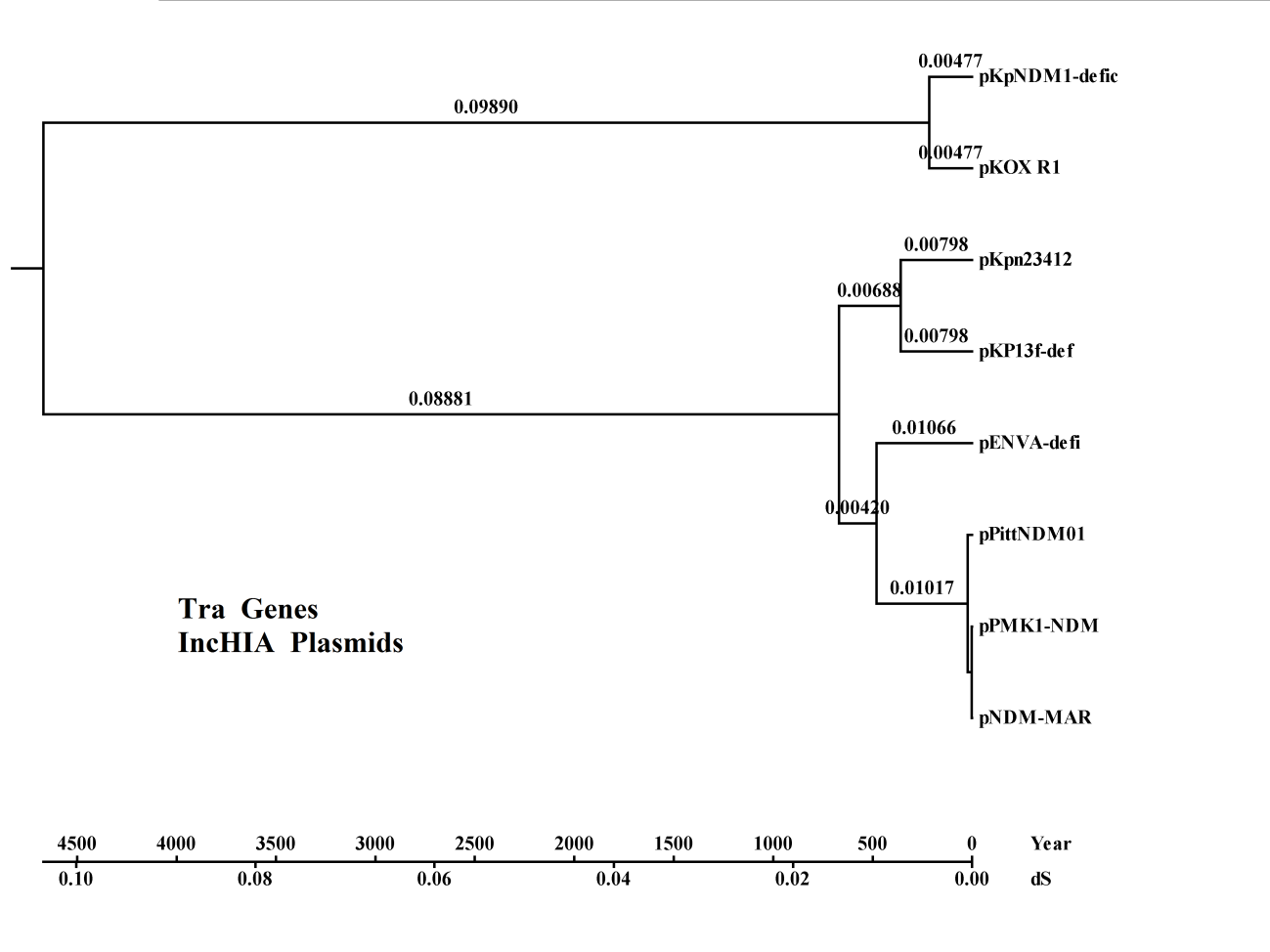

**Tra Genes Sub-Tree 4**.- IncHIA plasmids from the expanded FIGURE 1.
