## Supplementary material for "Comparative genomics and phylogeny of sequenced IncHI plasmids": Tellurite genes sub treee

**TELLURITE GENES SUB-TREES**

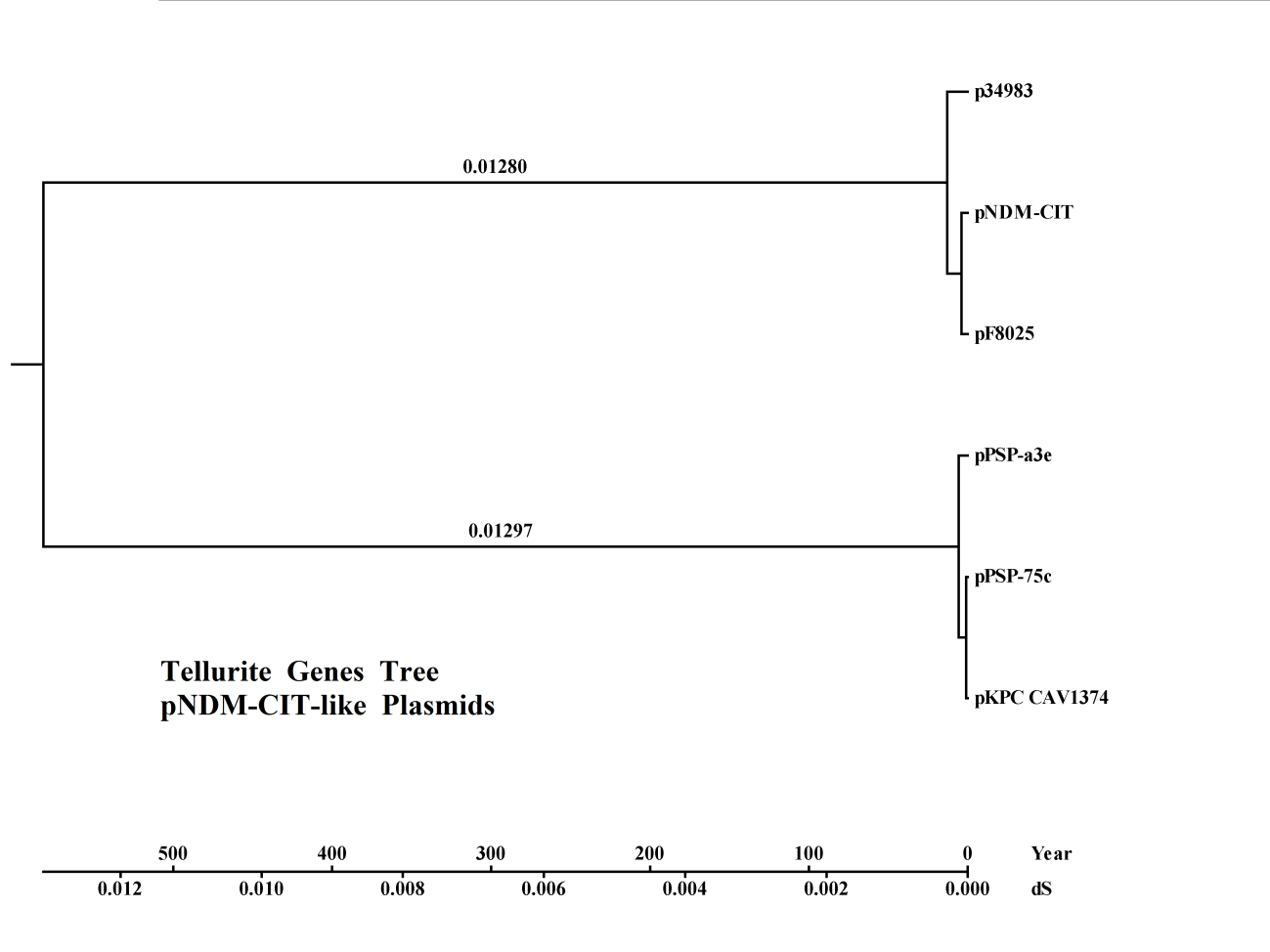

**Tellurite Genes Partial Tree 1 .-** IncHI pNDM-CIT-like Plasmids

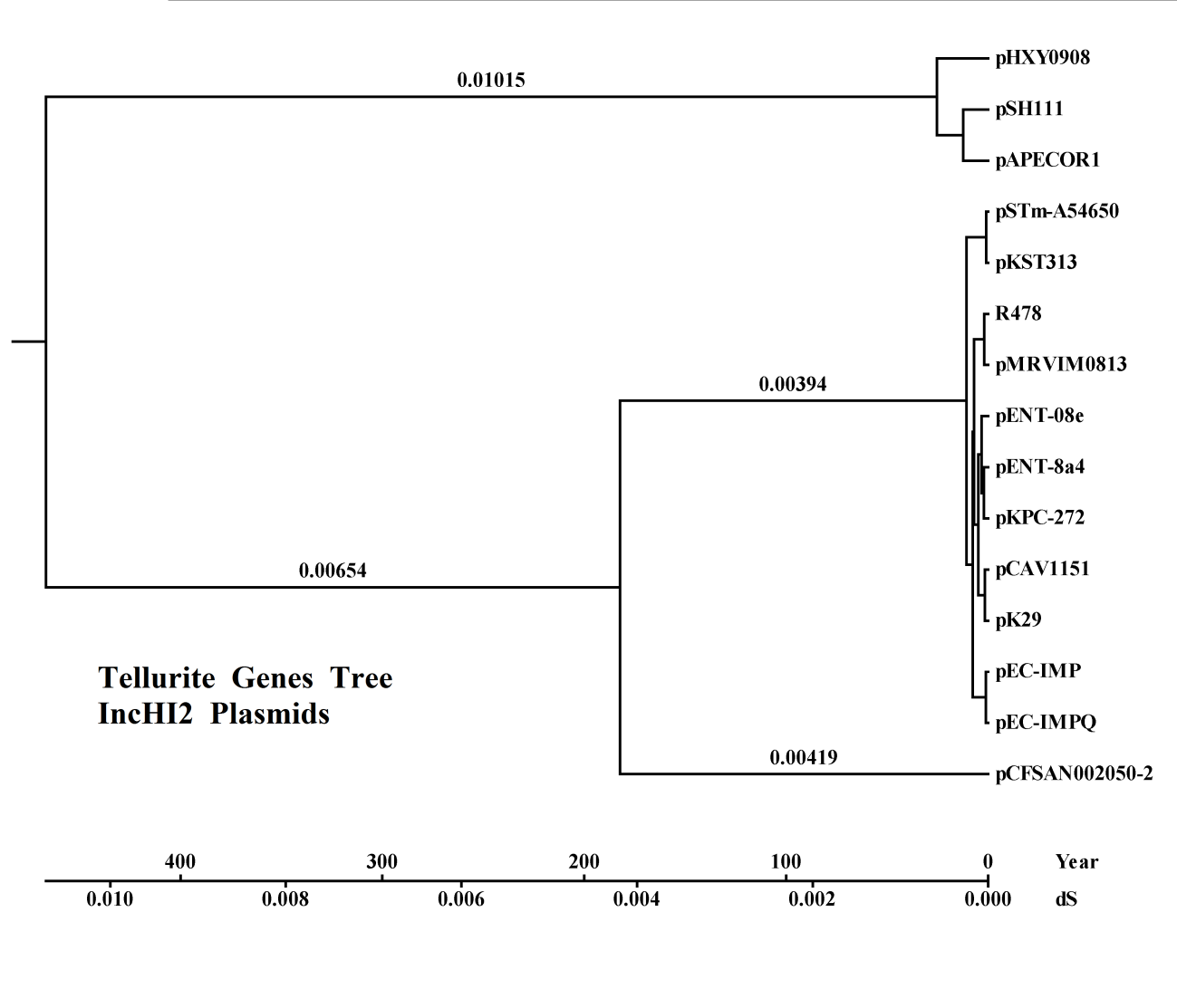

**Tellurite Genes Partial Tree 2.-** IncHI2 Plasmids.

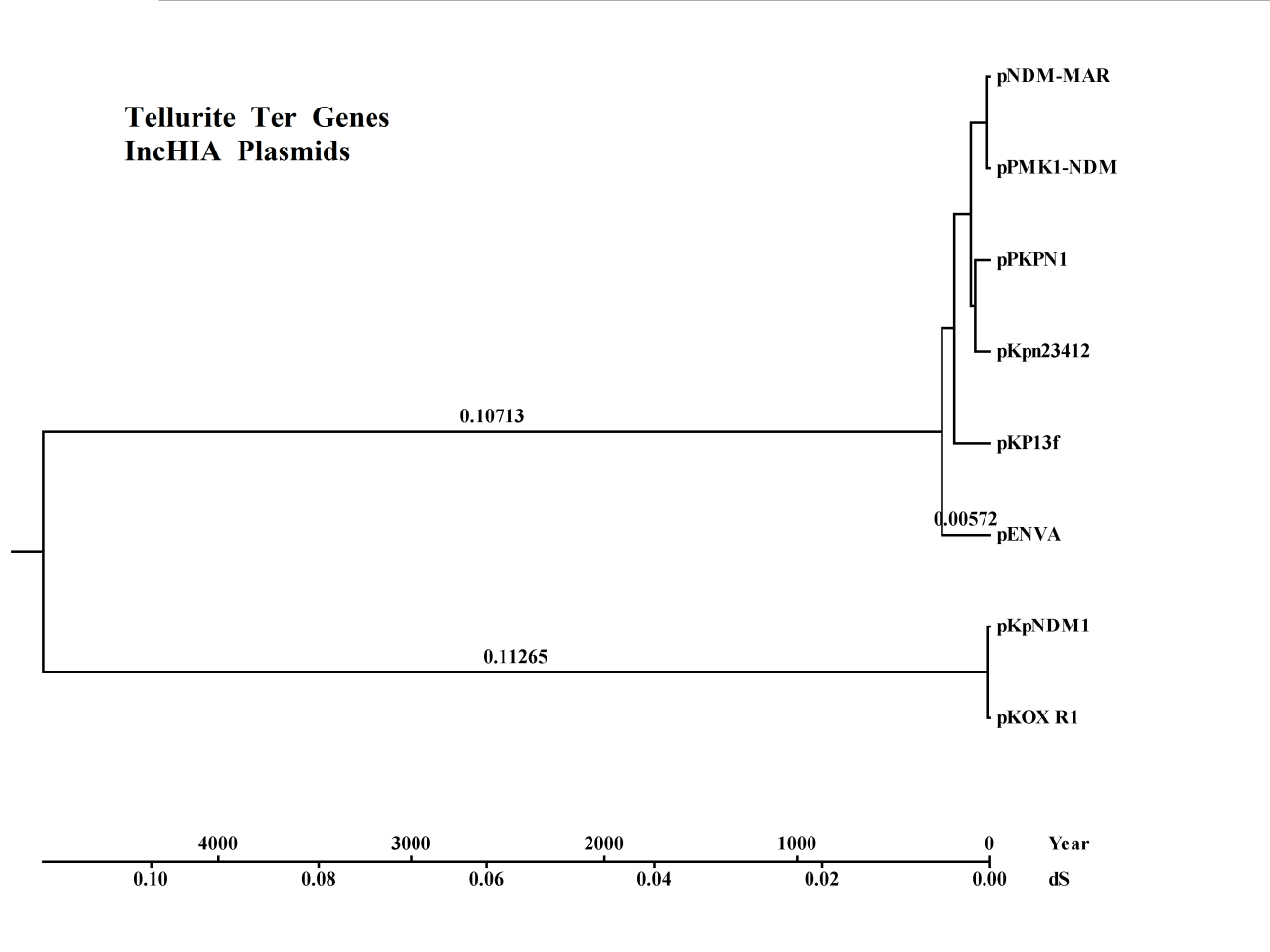

**Tellurite Genes Partial Tree 3.- IncHIA plasmids**
